# Tetraspanin CD82 reduces the formation of CADM1 oligomers

**DOI:** 10.64898/2025.12.12.693935

**Authors:** Elisa Lamottke, Harry Warner, Sjoerd van Deventer, Fabian Schwerdtfeger, Linde Lanting, Eric G. Huizinga, Annemiek B. van Spriel, Piet Gros

## Abstract

CD82 is a member of the tetraspanin protein superfamily and known as a metastasis suppressor. We identified cell adhesion molecule 1 (CADM1) as an interaction partner of CD82. CADM1 mediates cell adhesion by forming *cis* and *trans* oligomers that connect membranes. We show that CD82 reduces the formation of CADM1 oligomers when solubilized by detergent, on liposomes and in *cellulo* using Jurkat T-cells. Our data is consistent with a 1:1 complex of CD82 and CADM1 in *cis* that leaves the CADM1 *trans*-interaction site accessible. Cryo-electron microscopy of the CD82:CADM1 heterodimer suggests an interaction site between the large-extracellular loop of CD82 and an Ig-like domain of CADM1. Consistently, liposomes coupled with CADM1 ectodomain show reduced clustering when reconstituted with CD82. We hypothesize that CD82 may affect spacing of the transmembrane helices of CADM1, possibly by interacting with the extracellular Ig-like domains and hence disrupting CADM1 oligomerization between membranes.

## Introduction

Tetraspanins are a family of 33 four-transmembrane-helix proteins in humans, are widely expressed in multicellular organisms and play key roles in cell signaling, adhesion, and membrane organization [1, 2, 3]. They organize the cell membrane by interacting with other membrane proteins and in some cases regulate their surface expression [2, 4]. Super-resolution microscopy studies revealed that tetraspanins organize their partner protein in small clusters, called tetraspanin-enriched microdomains (TEMs) or nanodomains [3, 5, 6, 7]. Recent cryo-electron microscopy (cryo-EM) studies show different ways of organization of tetraspanins and their partner proteins. Some interaction partners appear as hetero-dimers or -tetramers [8, 9, 10, 11]. In the urinary tract, tetraspanins UPIa and UPIb form 16 nm complexes, consisting of six UPIa:UPII and six UPIb:UPIIIa hetero dimers that arrange into quasi-crystalline networks [12]. In the retina, the tetraspanins ROM1 and peripherin-2 form hetero-dimers that assemble into large oligomeric complexes [13, 14]. The interaction sites between tetraspanins and their partner proteins were observed to be in the extracellular domain [8, 9], the transmembrane domain [10, 11], or both [12]. Studies on CD19-CD81 fusion protein and CD45:CD53 complexes show that the interaction with partner proteins in TEMs might be affected by conformational changes of the tetraspanin [9, 15]. Tetraspanins can adopt an open conformation, where the transmembrane helices arrange in parallel and the large extracellular loop (LEL) extends from the membrane, or a closed conformation in which the helices adopt a cone-like shape with the LEL closing it off [9]. Switching between the open and closed conformation may be regulated by lipids, as might be the case for gangliosides and CD82 [16, 17] and has been suggested for cholesterol and CD81 [18, 9]. In the case of CD81, the crystal structure showed cholesterol bound between the transmembrane helices in a closed conformation [18] whereas the cryo-EM structure of the CD81-CD19 complex, showed CD81 in an open conformation without cholesterol [9].

Tetraspanin CD82 functions as a metastasis suppressor [19] as demonstrated by its ability to inhibit cell proliferation and adhesion through interactions with receptor tyrosine kinases and integrins in solid tumors [20]. The invasion of tumor cells into surrounding tissues is inhibited by CD82-mediated increase in cell adhesion, as shown for prostate cancer cells [21]. On the other hand, CD82 was also shown to decrease cell adhesion during tumor invasion by indirect inhibition of plasminogen that degrades the extracellular matrix and therefore facilitates invasion in new tissue [22]. CD82 consists of 267 amino acids and shows the typical features of the tetraspanin family, including four transmembrane helices, a small extracellular loop (SEL), and a large extracellular loop (LEL) with a CCG motif. The CD82 LEL consists of 118 amino acids and contains three N-glycosylation sites which make it relatively large compared to other tetraspanins [23].

Cell adhesion molecule 1 (CADM1) is a widely expressed adhesion molecule that is involved in several biological functions. CADM1 overexpression promotes recruitment of T cells or mast cells in autoinflammatory diseases, like type 1 diabetes, asthma or neuroinflammatory diseases [24, 25, 26]. In cancer, CADM1 can function as an oncogene as well as a tumor suppressor by regulating different signaling cascades, such as the Hippo pathway that promotes cell proliferation [27]. In T-cell lymphoma, cell adhesion by CADM1 leads to increased organ infiltration of tumors, which makes it a suitable therapeutic target and a biomarker [28, 29]. CADM1 is a member of the immunoglobulin (Ig) superfamily. The 398 amino-acid long protein consists of three extracellular Ig-like domains that are N-glycosylated at six sites, a 45 amino acid long O-glycosylated [30] and non-globular linker or stalk domain and a transmembrane helix connecting to the intracellular signaling domain. The intracellular domain contains a 4.1 binding motif, that interacts with actin-binding protein DAL-1 [31] and a PDZ-binding motif that interacts with scaffolding proteins in neurons and cancer cells [32, 33]. CADM1 can dimerize with itself, other members of the CADM family, and the cytotoxic and regulatory T-cell molecule (CRTAM) [34, 28, 35]. CADM1 dimerizes in *cis* and *trans* which leads to the formation of oligomers and consequential cell adhesion [34]. *Cis* dimers are thought to form via interactions of the second or third Ig domain [34, 36] and the transmembrane domain that contains a GxxxA motif, which is typical for receptor dimerization [37, 38]. The first Ig domain is responsible for forming dimers in *trans* [36, 39]. It is proposed that *cis* dimers are formed first and stabilize the formation of *trans* dimers, resulting in a so-called zipper motif in between membranes [40, 41].

In this study, we identified human CADM1 as an interaction partner of human tetraspanin CD82. To investigate the biological function, CADM1 oligomerization was analyzed in presence and absence of CD82 by biophysical assays and in *cellulo*. We further characterized the interaction site of CADM1 and CD82 by cryo-EM.

## Results

### Identification of CADM1 and CD82 as interaction partners

To identify new CD82 interaction partners, CD82, fused C-terminally with an eGFP-StrepII_3_-tag, was overexpressed in HEK-E+ cells, solubilized and purified via its StrepII_3_-tag in DDM. CD82-eGFP-StrepII_3_ appeared as a double band at 50 kDa and 70 kDa (which includes 30 kDa of eGFP-StrepII_3_) on SDS-PAGE gel, identified through the eGFP-signal (**Fig. S1A-B**). The double band was likely due to N-glycosylation variants, as often observed for tetraspanins and other N-glycosylated proteins [10, 42, 43]. Protein bands with molecular weights (MW) of approximately 90-100 kDa, 100-110 kDa and >250 kDa were cut from the SDS-PAGE gel and identified using mass spectrometry (**Fig. S1C**). CADM1 was among the ten proteins with the highest scores in the two lower MW samples and was the only membrane protein localized in the cell surface that co-purified with CD82 (**Table 1**). Other high-scoring proteins included molecular chaperones e.g. calnexin and heat-shock proteins, consistent with protein over-expression of membrane proteins [44, 45]. Next, we co-expressed CD82-StrepII_3_ and CADM1-His_6_ and solubilized and purified these using DDM with strep-tactin resin. Purification yielded bands observed at 30, 40, 70, 90 and 250 kDa in SDS-PAGE gel (**Fig. 1A**). The double band at 70 and 90 kDa was identified as CADM1-His_6_ via an anti-His western blot (**Fig. 1B**). The bands at 30 and 40 kDa were consistent with the double band at 50 and 70 kDa of CD82-eGFP-StrepII_3_ and therefore can be attributed to CD82-StrepII_3_; the 250 kDa band was likely an impurity. To confirm the formation of a complex we performed fluorescent size exclusion chromatography (F-SEC) monitoring the eGFP signal. For this we expressed CADM1-eGFP-StrepII_3_, CD82-eGFP-StrepII_3_ and co-expressed CD82-eGFP-StrepII_3_ and CADM1-His_6_. Purified CADM1-eGFP-StrepII_3_ appeared as a smeared band at ca. 100-110 kDa (compared to 70 and 90 kDa without the 30 kDa eGFP-StrepII_3_-tag) on SDS-PAGE gel and CD82-eGFP-StrepII_3_ at 50 and 70 kDa (**Fig. 1C**). Co-expression of CD82-eGFP-StrepII_3_ and CADM1-His_6_ resulted in the expected double band at 50 and 70 kDa for CD82-eGFP-StrepII_3_ but no band was observed at the expected MW of CADM1-His_6_ (**Fig. 1C**). However, co-purification of CADM1-His_6_ was confirmed by an anti-His western blot showing a high MW band of 90 kDa, consistent with the top band of 90 kDa in the previous co-purification of CD82-StrepII_3_ and CADM1-His_6_. No 70-kDa band of CADM1-His_6_ is visible, which is possibly due to reduced yield of the protein complex (**Fig. 1D, B**). On F-SEC, CADM1-eGFP-StrepII_3_ displayed a main peak at 10.5 mL with additional peaks between 8-9.5 mL and 12-14 mL (**Fig. 1E**). The peaks at multiple elution volumes (EV) might represent different oligomeric states. The F-SEC chromatogram of purified CD82-eGFP-StrepII_3_ showed a main peak at 12 mL, while co-purification of CD82-eGFP-StrepII_3_ and CADM1-His_6_ showed an additional peak at 10 mL, indicating the occurrence of a CADM1-His_6_, CD82-eGFP-StrepII_3_ complex. In contrast to CADM1-eGFP-StrepII_3_ alone, the complex of CD82-eGFP-StrepII_3_:CADM1-His_6_ eluted in one peak and showed no indication of CADM1 oligomerization.

**Figure 1:**
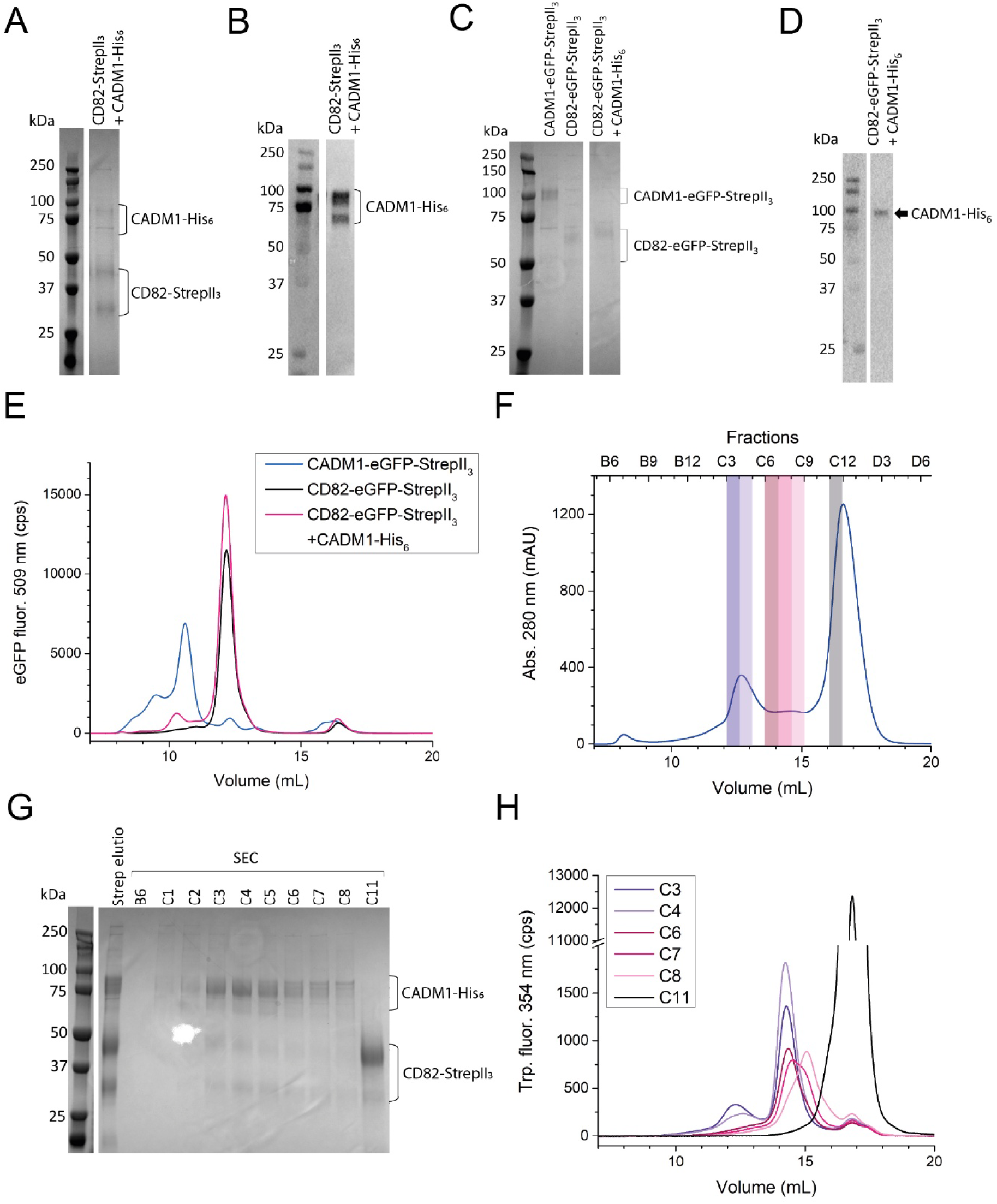
Co-purification of CADM1 and CD82. (A, B) 10% SDS-PAGE gel of co-expressed CADM1 and CD82 after purification using strep-tactin (A) stained with Coomassie and, (B) anti-His western blot. (C, D) 10% SDS-PAGE gel of strep-tactin-purified individually and co-expressed CADM1 and CD82 (C) stained with Coomassie and, (D) anti-His western blot of co-purified CD82 and CADM1. (E) F-SEC of samples shown in C visualizing the eGFP signal using a Superdex 200 increase 10/300 GL column (F) SEC-profile of strep-tactin-purified CD82 and CADM1 large-scale expression using a Superose 6 increase 10/300 column. Single fractions were analyzed by (G) SDS-PAGE gel stained with Coomassie and fractions marked with transparent, colored boxes (H) were reinjected onto the same column and monitored via the tryptophan fluorescence signal.

**Table 1:**
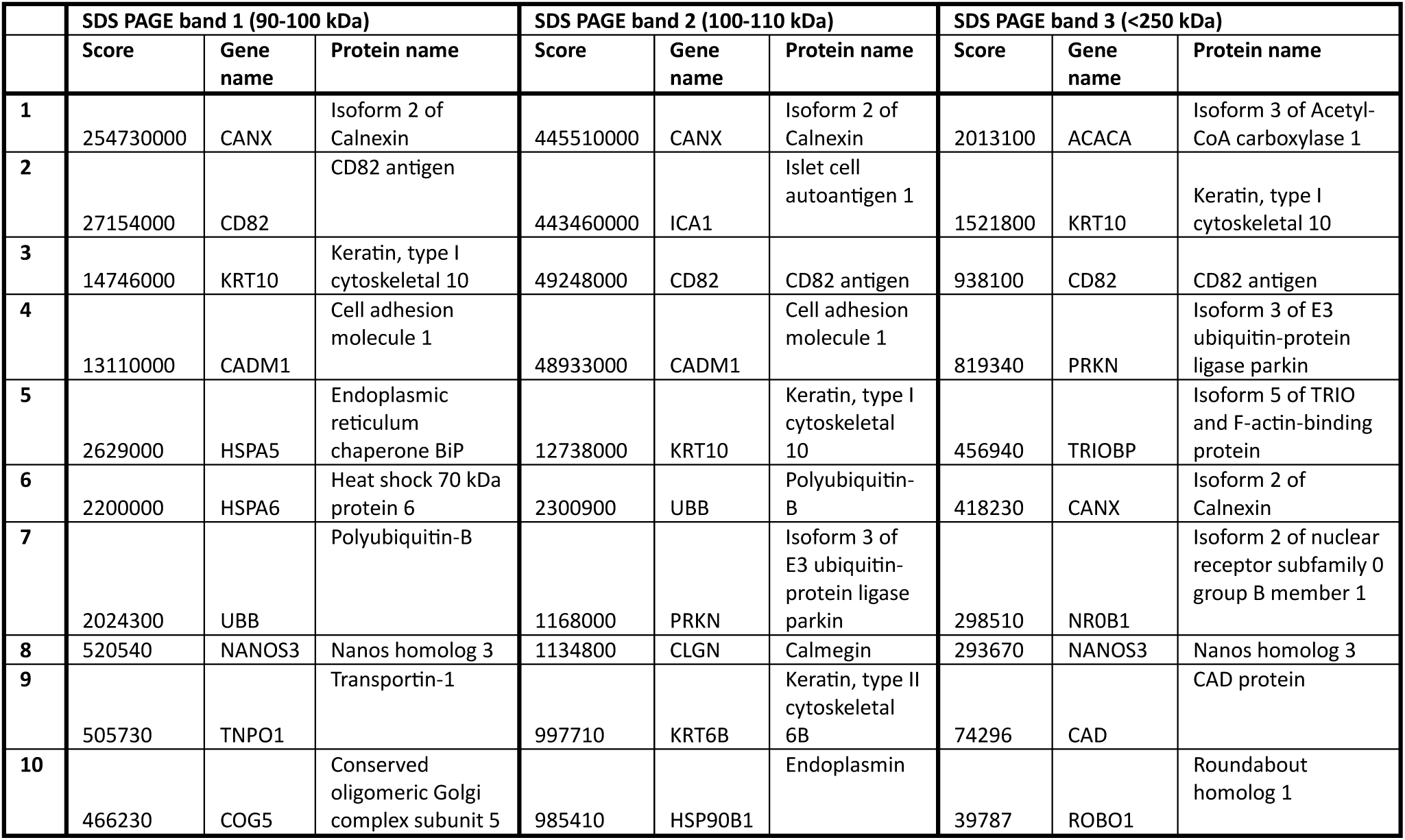
Highest scores from mass spectrometry. Top iBAQ scores detected in mass spectrometry of each analyzed SDS-PAGE band containing proteins that were co-purified with CD82-StrepII_3_ from HEK-E+ cells (**Fig. S1**).

We co-expressed CD82-StrepII_3_ and CADM1-His_6_ in larger culture volume and used strep-tactin and SEC for protein purification in DDM. The SEC profile qualitatively resembled the SEC profile of the small-scale co-expression displaying a high peak at 17 mL, a lower peak at 12.5 mL with a non-zero intensity region in between (**Fig. 1F**). The peak at 17 mL contained only CD82-StrepII_3_, visible as a double band at 30 kDa and 40 kDa, while all fractions between 16 mL and 12.5 mL EV contained both, CD82-StrepII_3_ and CADM1-His_6_ (**Fig. 1G**). The latter showed multiple bands between 70-90 kDa in SDS-PAGE gel, consistent with the anti-His western blot of the small-scale purification without an eGFP-tag (**Fig. 1A-B, G**). Fractions containing CD82-StrepII_3_:CADM1-His_6_ complex showed the 30 and 40-kDa bands at approximately equal intensity, whereas the peak containing only CD82-StrepII_3_ displayed a significantly more intense 40 kDa band. Using the same SEC column used for purification, we analyzed selected fractions for stability of potential complexes (**Fig. 1H**). Diluted complex, in fractions C3 and C4 from the 12.5 mL peak, eluted partly at 12.5 mL and mostly in a defined peak at 14 mL, suggesting the existence of a weakly interacting higher-order complex eluting at 12.5 mL. Reinjected fractions C6-C8, corresponding to the broad region between 13.5 mL and 15 mL EV, eluted at their approximate original volume. However, consecutive fractions showed a gradual change in elution profile suggesting the presence of two populations, one eluting at 14 mL, the other at 15 mL, possibly caused by dissociation of the micellar complex during SEC. All reinjected fractions showed a small peak at the EV of CD82-StrepII_3_ indicating some CD82 dissociated from complexes originally present in these samples. Fraction C11 eluted as a single peak at the EV of CD82-StrepII_3_ confirming it contains pure protein.

In conclusion, we showed that CADM1 solubilizes and co-purifies with CD82. CADM1-eGFP-StrepII_3_ alone eluted as multiple peaks in SEC, most likely representing different oligomers, while CADM1-His_6_:CD82-StrepII_3_ complex eluted as two distinct, a low and a high-order, complexes.

### Characterization of CADM1-CD82 complexes

We expressed CADM1 and CD82 fused by a 31 amino-acid linker (4xGGS-TEV site-4xGGS), CADM1-CD82-StrepII_3_, and compared it to CADM1-StrepII_3_ by F-SEC monitoring tryptophan fluorescence (**Fig. S2A-D, 2A-B**). CADM1-StrepII_3_ showed a peak at 14 mL and a shoulder at 12.5 mL for the two lowest concentrations used, 0.15 µM and 0.7 µM (**Fig. 2A**). At 7 µM, the main peak and shoulder shifted to smaller EVs, while the height of the shoulder increased and shifted to 12 mL, indicating the formation of higher order CADM1-complexes. Concentrations of 68 µM and 80 µM showed an even further shift of the main peak to 12 mL with a prominent shoulder at 10 mL and a smaller shoulder at 14 mL (**Fig. 2A**). Similar profiles, though shifted due to the fused CD82 molecule, were observed at the lowest concentrations (0.15 µM-7 µM) used for CADM1-CD82-StrepII_3_ (**Fig. 2B**). At high CADM1-CD82-StrepII_3_ concentrations (68 µM and 80 µM), peaks at 14 and 12 mL remained, but the shoulder at 10 mL was markedly reduced compared to CADM1-StrepII_3_ (**Fig. 2B**). This trend appeared to continue at higher concentrations as indicated by the SEC profiles obtained during protein purification of CADM1-StrepII_3_ and CADM1-CD82-StrepII_3_ (**Fig. S2A, S2C**). Finally, all chromatograms showed a peak at 16 mL that may be attributed to free CD82-StrepII_3_ (**Fig. 2B, S2C-D**). Overall, the EVs of different complexes were consistent between CADM1-CD82-StrepII_3_ fusion protein and non-fused CD82-StrepII_3_ and CADM1-His_6_ (**Fig. 2B, 1E**). Compared to CADM1 alone, CADM1-CD82 showed reduced intensity of the earliest eluting peak.

**Figure 2:**
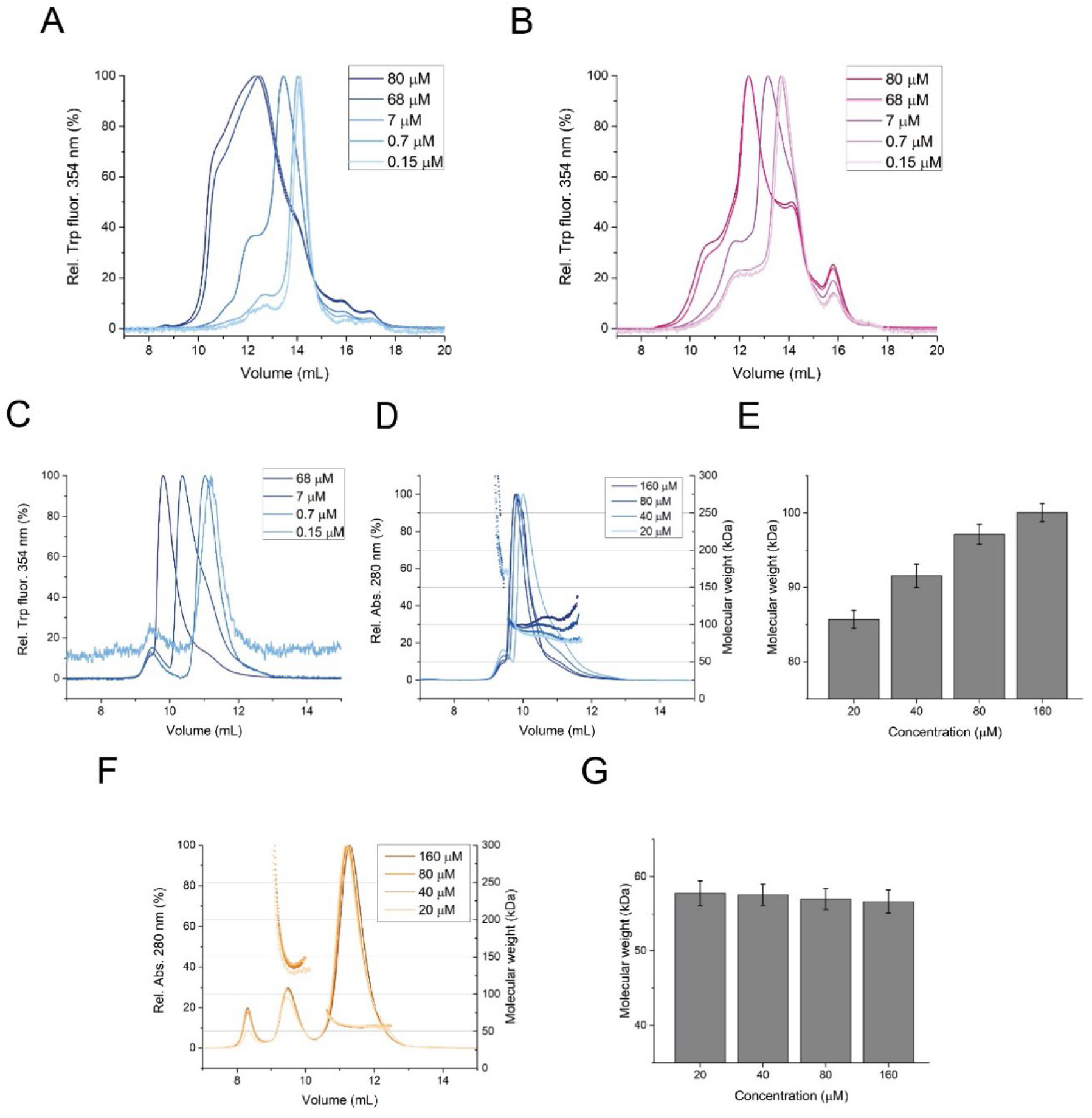
Characterization of CADM1 oligomerization and CADM1-CD82 complex formation. (A) Tryptophan fluorescence elution profile of different concentrations of purified CADM1-StrepII_3_ and, (B), CADM1-CD82-StrepII_3_ fusion protein injected onto a Superose 6 increase 10/300 column. (C) F-SEC of different concentrations of CADM1(ECD)-His6 using a Superdex 200 increase 10/300 column. (D) SEC-MALLS analysis of CADM1(ECD)-His_6_ at different concentrations under the same conditions as in C (E) Molecular weight determined from the main peak in panel D eluting at a volume of 10 mL for each concentration. (F) SEC-MALLS analysis of different concentrations of CADM1(ECD,F42S)-His_6_ using a Superdex 200 increase 10/300 column. (G) Molecular weight determined from the main peak in panel F eluting close to 11.2 mL.

To gain further insight into the oligomerization, we expressed and purified His-tagged CADM1 ectodomain, CADM1(ECD)-His_6_ (**Fig. S3A-B**). In analytical F-SEC of CADM1(ECD)-His_6_ (**Fig. 2C**), the main peak gradually shifted from 11.5 to 10 mL with increasing concentrations and showed tailing towards high EVs at high concentrations. This gradual shift was consistent with the shift from 14 mL to 12 mL of full-length CADM1-StrepII_3_. However, no early eluting peak or shoulder was observed for CADM1(ECD)-His_6_. Furthermore, we performed SEC combined with multiple angle laser-light scattering (SEC-MALLS) at concentrations of 160 µM, 80 µM, 40 µM and 20 µM, all resulting in a similar SEC profile as CADM1(ECD)-His_6_ at 68 µM (**Fig. 2D**). The MALLS signal resulted in calculated MWs ranging from 85 kDa to 100 kDa, which is in between the expected weight of a 60 kDa monomer and a 120 kDa dimer (**Fig. 2E**). Hence, CADM1(ECD)-His_6_ appears as a monomer-dimer equilibrium on SEC, with the dimer eluting close to 10 mL and the monomer eluting close to 11.5 mL (**Fig. 2C-D**). To determine if the observed dimer is in *cis* or *trans* configuration, we repeated SEC-MALLS with a F42S mutant of CADM1, (CADM1(ECD,F42S)-His_6_), that blocks the formation of *trans*-dimers [28] (**Fig. S3C-D**). The largest peak was observed at 11.5 mL for all injected concentrations, similar to CADM1(ECD)-His_6_ at low concentration (**Fig. 2F**). MALLS indicated an MW of 57 kDa, which is consistent with the expected 40 kDa of monomeric CADM1(ECD) plus six N-glycans of approximately 3 kDa each (**Fig. 2G**). Therefore, the observed fast-exchange monomer-dimer equilibrium is likely caused by the *trans*-dimer formation of CADM1. CADM1(ECD,F42S)-His_6_ and CADM1(ECD)-His_6_ both showed an additional peak at 9.5 mL, consistent with the SEC profile of CADM1(ECD,F42S)-His_6_ during protein purification (**Fig. S3C**). The SEC profile of CADM1(ECD)-His_6_ might contain a similar peak overlapping with the main peak (**Fig. S3A**). For CADM1(ECD,F42S)-His_6_, MALLS determined an MW of ca. 140 kDa for this peak at all concentrations (**Fig. 2F**), while the MW of the corresponding peak for CADM1(ECD)-His_6_ could not be determined due to overlap with the dimer peak (**Fig. 2C**). The EV of the peak at 9.5 mL did not change with protein concentration as would be expected for a potential dimer-trimer equilibrium. Thus, this peak could represent misfolded protein, contamination or an additional CADM1 molecule interacting in *cis*.

With this information for soluble CADM1(ECD)-His_6_, we can now better interpret the SEC profiles of DDM-solubilized (full-length) CADM1-StrepII_3_ and CADM1-CD82-StrepII_3_ (**Fig. 2A-B**). At high concentrations, a higher-order oligomer eluted at 10 mL, which was reduced in the presence of CD82. This oligomer did not form with CADM1(ECD)-His_6_ and therefore possibly involves *cis* interactions, which have been reported to require the transmembrane helix of CADM1 [38]. At low concentrations both proteins showed a monomer-dimer equilibrium in *trans* configuration, like CADM1(ECD)-His_6_. To analyse the interaction between CADM1 and CD82, purified CADM1(ECD)-His_6_ was added to a final concentration of 0.1 mg/mL (2.5 µM) to cleared lysate of HEK-E+ cells expressing CD82-StrepII_3_ followed by purification on strep-tactin-resin (**Fig. S4A**). An anti-His western blot did not show co-purification of CADM1(ECD)-His_6_ at detectable amounts (**Fig. S4B**). This indicates that an interaction between the transmembrane helices of the two proteins is required for binding or might be a result of CADM1(ECD) not being inserted into a two-dimensional cell membrane or micelle. In conclusion, we attribute the elution at 14 mL to monomeric CADM1-CD82-StrepII_3_ fusion protein and the corresponding peak of non-fused CADM1-His_6_:CD82-StrepII_3_ to hetero-dimer in *cis* (**Fig. 2B,1F**). Accordingly, the complex eluting at 12 mL corresponds to a *trans* complex of the fused monomer or dimerization of the non-fused hetero dimers in *trans*.

### Effect of CD82 on clustering of CADM1-coated liposomes

We investigated how CD82 impacts CADM1-mediated cell adhesion using liposomes as a model. Extruded DOPC:DGS-NTA liposomes were incubated with different concentrations of CADM1(ECD)-His_6_ and liposome-cluster formation was analyzed using cryo-EM and dynamic light scattering (DLS). Liposomes coated with 20 µM CADM1(ECD)-His_6_ showed clusters of liposomes with a dense arrangement of protein in between adjacent membranes and excess CADM1(ECD)-His_6_ next to the clusters (**Fig. 3A**). Liposomes visibly aligned with an apparent distance of approximately 35 nm between membranes at several spots imaged at the edge of liposome clusters. The observed distance is consistent with a CADM1 *trans*-dimer model with three extended Ig-like domains with the terminal domain in overlap (five times ca. 3-4 nm) and a mostly extended conformation of the O-glycosylated linkers (two times ca. 9-10 nm) (**Fig. 3B**) [39, 35]. At lower magnifications of 7300x, the dense arrangement of liposomes covered with CADM1(ECD)-His_6_ was visible (**Fig. S5**). When covering liposomes with CADM1(ECD,F42S)-His_6_ less dense clusters were visible and when covering liposomes with EGF-His_6_ no clusters were observed (**Fig. S5**). This suggests non-complete disruption of CADM1 interactions between membranes by the F42S mutation and no interaction of liposomes covered with monomeric protein.

**Figure 3:**
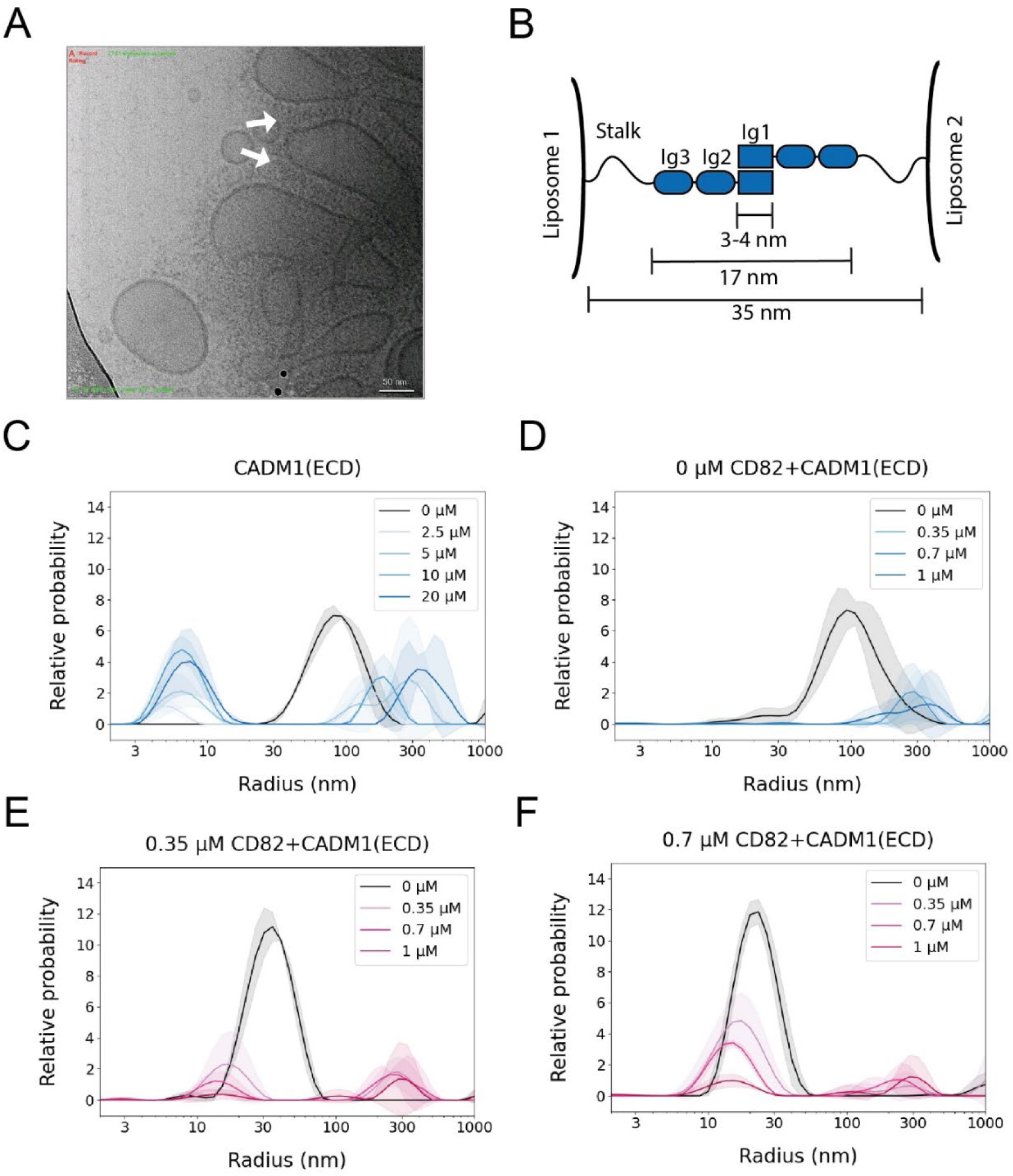
Liposome clustering by immobilized CADM1(ECD)-His_6_ in the presence and absence of CD82-StrepII_3_. (A) Cryo-EM image at 100,000 x magnification of DOPC:DGS-NTA (8:2) liposomes connected by CADM1(ECD)-His_6_ recorded on a Talos Arctica electron microscope. Liposomes were extruded through a 200 nm filter. White arrows indicate dense arrangements of CADM1(ECD)-His_6_ in between membranes. (B) Cartoon of CADM1(ECD) *trans* dimer between liposomes with indicated sizes of an average Ig-like domain, resulting size of Ig-like domains in *trans* conformation and observed distance between membranes in A. (C-F) DLS by DOPC:DGS-NTA (8:2) liposomes coated with CADM1(ECD)-His_6_. All DLS plots are cropped from 2 nm to 1,000 nm radius. Translucent areas represent the standard deviation of the relative probability. (C) DLS by liposomes diluted 10x compared to panel A extruded through an 800 nm filter and coated with different concentrations of CADM1(ECD)-His_6_ as indicated. (D-F) DLS by liposomes diluted 5x compared to panel A extruded through a 200 nm filter. Liposomes in (C) underwent reconstitution without inserting membrane protein in the membrane. Liposomes in (E,F) were reconstituted with different concentrations of CD82 as indicated.

Quantification of liposome size by DLS resulted in an average radius of 100 nm for uncoated liposomes (**Fig. 3C**). Adding CADM1(ECD)-His_6_ at concentrations of 2.5 µM, 5 µM, 10 µM, and 20 µM, resulted in sizes between 100 nm and 500 nm with high standard deviations, likely representing clusters of liposomes connected via CADM1(ECD)-His_6_ oligomers in *trans* (**Fig. 3C**). In addition to the clusters, we observed a peak at 6 nm, due to excess unbound CADM1(ECD)-His_6_ (**Fig. 3C**). Liposomes reconstituted with 0.35 µM or 0.7 µM CD82-StrepII_3_ were incubated with CADM1(ECD)-His_6_ at concentrations of 0.35 µM, 0.7 µM and 1 µM to achieve approximate equimolar concentrations with respect to reconstituted CD82-StrepII_3_. No excess of CADM1(ECD)-His_6_ was detected at this range of concentrations (**Fig. 3C-E**). The reconstitution procedure did not affect liposomes, as those reconstituted without inserting CD82-StrepII_3_ showed comparable sizes in DLS (**Fig. 3D**). When liposomes were reconstituted with freshly purified CD82-StrepII_3_, the liposomes displayed reduced radii of 40 nm (**Fig. S6A-B**), which was apparently due to insertion of membrane proteins, since this was also observed after reconstitution of an EGFR fragment, truncated after the transmembrane (TM) domain, EGFR(I-TM) (**Fig. 3D-E, S6C-D, S7A**). Consistent with previous experiments, the addition of CADM1(ECD)-His_6_ to CD82-resconstituted liposomes resulted in liposome clusters ranging in size from 100-500 nm radius. However, a subpopulation of liposomes with a radius between 20 nm and 30 nm did not cluster (**Fig. 3D-E**). The relative probability of this subpopulation increased when higher concentrations of CD82-StrepII_3_ were reconstituted and relative probability and observed sizes of liposomes decreased with the added concentration of CADM1(ECD)-His_6_. As we showed that reconstitution reduces the radius of liposomes, the smaller liposomes likely contain a higher concentration of CD82-StrepII_3_, thus indicating that reconstitution of CD82-StrepII_3_ prevents CADM1(ECD)-His_6_-induced liposome clustering. The observed trends were reproducible with freshly purified CD82 reconstituted in freshly extruded liposomes (**Fig. S8**). Liposomes reconstituted with EGFR(I-TM)-StrepII_3_ showed no cluster-resistant liposomes upon addition of CADM1(ECD)-His_6_ (**Fig. S7A**), showing that this is specific to CD82-reconstitutions. To confirm that liposome clustering is induced by CADM1(ECD)-His_6_ *trans-*interactions, CD82-StrepII_3_ reconstituted liposomes were incubated with the CADM1(ECD,F42S)-His_6_ mutant which resulted in no apparent clustering of liposomes (**Fig. S7B**). In conclusion, we showed that CADM1-mediated cell adhesion can be mimicked by anchoring CADM1(ECD)-His_6_ to DOPC:DGS-NTA (8:2) liposomes which is inhibited by reconstitution of CD82.

### CD82 inhibits CADM1 oligomerization in the plasma membrane of Jurkat cells

To test the impact of CD82 on CADM1 clustering in *cellulo*, we generated CD82 knockout (KO) Jurkat T-cells using CRISPR-Cas9 technology. We observed that genetic ablation of CD82 did not affect CADM1 surface expression (**Fig. 4A-B**). We then utilized Airyscan super-resolution microscopy to examine the distribution of CADM1 on the surface of WT and CD82KO Jurkat T-cells (**Fig. 4C**). Analysis of the co-localization of CADM1 with CD82 in WT Jurkat cells revealed consistent, partial co-localization (**Fig. 4C-E**), indicating that the regulation of CADM1 by CD82 may be context specific and/or spatial-temporally dynamic.

**Figure 4:**
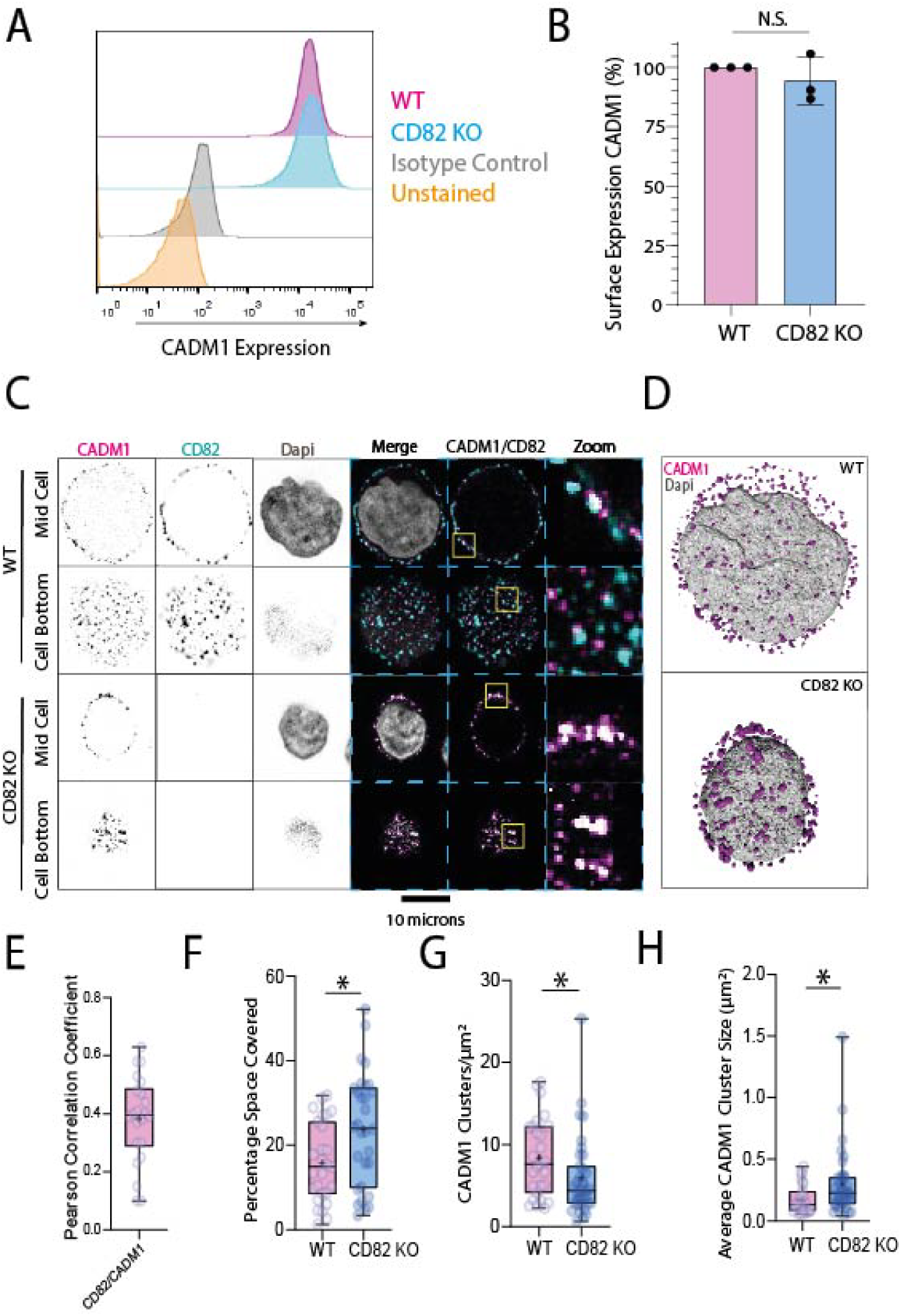
CD82 inhibits CADM1 clustering in *cellulo.* (A) Example flow cytometry histogram of the surface expression of CADM1 in WT and CD82 KO Jurkat cells. (B) Quantification of surface expression of CADM1 in WT and CD82 KO Jurkat cells of 3 independent experiments. (C) Airyscan microscopy of Jurkat cells (WT and CD82 KO) stained for CADM1, CD82 and DAPI (nucleus). (D) 3D rendering of cells from panel C. (E) Pearson’s correlation coefficient of CD82 with CADM1 in WT Jurkat cells (average of 27 cells over 3 independent experiments). (F) Quantification of the percentage space covered by CADM1 clusters in WT and CD82 KO Jurkat cells (minimum of 27 cells per condition over 3 independent experiments). (G) Quantification of the number of CADM1 clusters per unit area in WT and CD82 KO Jurkat cells (minimum of 28 cells over 3 independent experiments). (H) Quantification of average CADM1 cluster size in WT and CD82 KO Jurkat cells (minimum of 27 cells over 3 independent experiments). Statistical significance was calculated using two-tailed unpaired t tests; ∗p < 0.05. For box-and-whisker plots, box represents 25th to 75th percentile, whiskers represent maximum and minimum values, middle band represents data median, and + represents data mean.

To analyze the effect of CD82 knockout (KO), we first analyzed the percentage surface space occupied by CADM1. This revealed that upon loss of CD82, CADM1 occupies a greater percentage of the cell surface, compared to CADM1 on WT Jurkat cells (**Fig. 4F**). We then analyzed the number of CADM1 clusters, which revealed a reduction in the number of CADM1 clusters in CD82 KO Jurkat cells compared to WT cells (**Fig. 4G**). Following this, we analyzed the average size of CADM1 clusters. This revealed that loss of CD82 resulted in larger CADM1 clusters on average (**Fig. 4H**). Taken together, these results demonstrate that CD82 limits CADM1 nanodomain clustering at the plasma membrane, restricting their size. Conversely, the genetic loss of CD82 enables CADM1 clusters to form higher order clusters. This results in a smaller number of larger CADM1 nanodomains at the plasma membrane. Thus, tetraspanin CD82 restricts the clustering of CADM1 both in *vitro* and in *cellulo*.

### Characterization of CADM1-CD82 interaction site

We used single particle cryo-EM to further characterize the interaction between CADM1 and CD82. The complex of CADM1-His_6_ and CD82-StrepII_3_ was purified (**Fig. S9A-B**) and two samples, one from the 12 mL EV peak (fractions E3-E7), likely representing a 2:2 hetero-complex in *trans*, and one from the 14 mL EV peak (fractions E9-F1), likely representing a 1:1 hetero-complex, were plunge frozen. We collected a data set of 1,381 movies of the 14 mL sample on a Talos Arctica electron microscope (200 kV). The resulting 2D classes showed micelles with a protein density visible within and an elongated density sticking out of them, possibly representing a part of CADM1. Other 2D classes displayed connected or disconnected micelles in close proximity (**Fig. 5A**). Overall, the 2D classes were blurry and CADM1 was not visible in full-length, indicating flexibility. A second data set of 8,706 movies was collected of the 12 mL peak on a Krios electron microscope (300 kV). The resulting 2D classes showed similar particles to the 14 mL data set and in addition, 2D classes showing an elongated protein density without micelles (**Fig. 5B**). These additional 2D classes possibly represented a *trans*-dimer of CADM1:CD82 hetero complexes, which would be consistent with two micelles connected by an elongated protein density observed in the micrographs and our SEC data (**Fig. S10, 2**). Due to the high flexibility of the *trans*-dimer, we did not obtain 2D classes in full-length or a high-resolution density in 3D. 3D reconstruction from particles in 2D classes showing a single micelle with protein density inside resulted in a density map at a resolution of 9 Å (**Fig. 5C**). The density showed a micelle with additional density on top, that is very narrow close to the micelle and more globular shaped when further away. The handedness of the map cannot be determined with certainty due to low resolution (**Fig. S11A, B**). The density inside the micelle was consistent with multiple TM helices (**Fig. 5D**). AlphaFold3 predicted a CD82:CADM1 interaction between hydrophobic transmembrane helices and the LEL of CD82 and one of the three Ig-like domains of CADM1 in *cis*, with different models predicting an interaction with different Ig like domains (**Fig. 5E, S12**). All models had low pTM scores of 0.41 and 0.42 and three out of five predicted an interaction of Ig-like domain 3 with the LEL of CD82 with high local pLDDT values and expected position error of 15-20 Å (**Fig. 5E, S12**) [46, 47]. In conclusion, our cryo-EM data suggests an interaction between CD82(LEL) and an Ig-like domain of CADM1.

**Figure 5:**
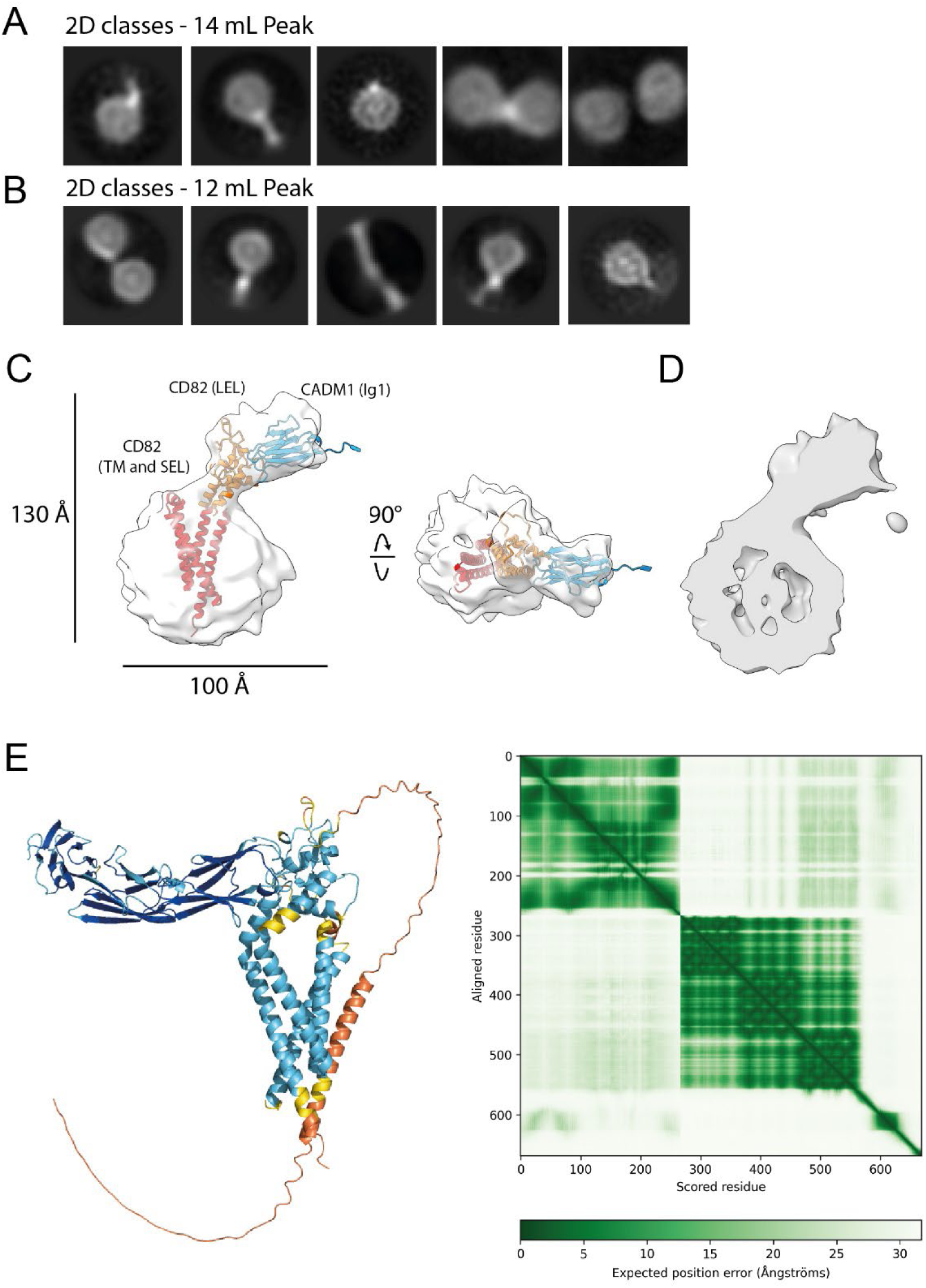
Structural evaluation of CADM1-CD82 interaction site. (A, B) Representative 2D classes resulting from data collection on 14 mL (A) and 12 mL (B) gel filtration peaks of CADM1-His_6_ and CD82-StrepII_3_ (Fig. S10). (C) 3D volume resulting from data collection of 12 mL peak manually fitted with rigid body models of CD82 transmembrane helices and SEL (red), CD82(LEL) (orange) and CADM1(Ig1) (blue) obtained from an AlphaFold3 prediction. (D) Cut-through of the 3D volume shown in panel C. (E) One out of five AlphaFold3 models CD82 and CADM1 colored by pLDDT score and predicted alignment error [46, 47].

## Discussion

In this study, we identified CADM1 as an interaction partner of tetraspanin CD82 by mass spectrometry. We confirmed the interaction by co-purification of CADM1 and CD82 and co-elution of both proteins in SEC. Analysis of individual SEC fractions by SDS-PAGE gel after large scale purification showed pure proteins supporting the conclusion that the interaction between CD82 and CADM1 is direct and does not involve additional proteins. CADM1 is a member of the immunoglobulin superfamily, which is common to several interaction partners of tetraspanins, such as EWI2, CD19 and N-cadherin [48, 49, 50].

CD82 was found to reduce the formation of CADM1 oligomers, which might result in an inhibiting effect on CADM1-mediated cell adhesion. Our analytical SEC data showed decreased oligomerization in the presence of CD82. Likewise, a subpopulation of CADM1(ECD)-coated DOPC:DGS-NTA liposomes resisted clustering in the presence of CD82. This subpopulation increased with the amount of reconstituted CD82 and decreased with increasing concentrations of CADM1(ECD). The average radius of liposomes decreased upon reconstitution of CD82 or EGFR(I-TM). Additionally, the radius of the cluster-resistant subpopulation decreased when adding higher CADM1(ECD) concentrations. We speculate that smaller liposomes contained higher concentrations of CD82 and were therefore more resistant to CADM1-induced clustering. Consistent with our in *vitro* assays, knock out of CD82 in Jurkat cells resulted in decreased numbers and increased size of CADM1 clusters, suggesting that the effect of CD82 is relevant in biological systems. Although large, TEM-like clusters were detected on Jurkat cells, we purified relatively small complexes. Possibly, additional cellular factors or weak TEM interactions are lost during solubilization and purification, resulting in small complexes between tetraspanins and partner proteins [51, 8, 9, 10, 11]. Taken together, we observed a moderate, but consistent inhibitory effect of CD82 on the formation of CADM1 oligomers in three different assays, representing increasing levels of physiological relevance. Nevertheless, it remains to be determined whether CD82 affects CADM1 function, since the contribution of CADM1 to Jurkat T-cell adhesion was reported to be limited [27] which may be explained by the expression of multiple different adhesion receptors on T cells. CADM1 expression in adult T-cell leukemia and breast cancer is associated with more aggressive organ infiltration [29, 52, 28], whereas CD82 is a known metastasis suppressor of solid tumors [19]. Accordingly, CD82 expression was shown to correlate with decreased survival rate in breast cancer cells [53], but has not been linked to adult T-cell leukemia. In another hematological cancer, acute amyloid leukemia, clustering of a closely related adhesion molecule, N-cadherin, was shown to be regulated by N-glycosylation and palmitoylation of CD82 [50]. Consistent with this observation, our interaction studies suggest an effect of N-glycosylation on CADM1:CD82 complex formation and N-glycosylation stabilizes the CADM1 dimer in *trans* [39]. Future research should address whether CD82 inhibits CADM1-driven tumor invasion and whether this effect is modulated by post-translation modifications.

We further investigated the potential mechanism by which CADM1 oligomerization is inhibited. SEC of CADM1:CD82 complex revealed the existence of at least two distinct complexes. F-SEC on CADM1 and CADM1 fused to CD82 combined with SEC-MALLS on CADM1(ECD) implied that CD82 and CADM1 form a 1:1 complex in *cis* that can dimerize to form a hetero-tetramer by CADM1 interaction in *trans*. In our SEC experiments, CADM1 dimerization in *trans* occurred at lower concentrations than oligomerization by *cis* interactions. This is in apparent contrast to a previous proposed model, that suggests CADM1 first forms dimers in *cis* and then further oligomerizes in *trans* [40, 41]. This contradiction can be explained by the use of detergent solubilized proteins in our studies. CADM1 is no longer restricted to a two-dimensional membrane, which would reduce the formation of dimers in *cis* configuration. In corroboration, the *cis* interaction was not observed for CADM1(ECD) in solution, however previous research has shown that the CADM1(ECD) is sufficient for cis dimerization when attached to the membrane via a GPI-anchor [40]. This is in agreement with observed particles on plunge frozen grids, that likely displayed *trans* interactions only for highly concentrated sample containing a 2:2 tetramer.

CADM1 and CD82 likely interact with each other through the CD82-LEL and an Ig-like domain of CADM1, as indicated by our low-resolution cryo-EM data. Our liposome-assay set up supports this notion, since reduced liposome clustering was observed when using liposomes with reconstituted CD82 and surface-coupled ectodomain of CADM1. In contrast, our studies showed no interaction between CD82 and CADM1(ECD), suggesting membrane helices are involved or a lack of interaction caused by reduced affinity when both protein partners are not confined to the same membrane. CD82 induced reduction of CADM1 oligomerization, liposome clustering and CADM1-cluster size on Jurkat T-cells can be explained by two mechanisms: i) CD82 causes steric hindrance of *cis*-interacting CADM1 molecules, possibly through interacting ectodomains; or ii) interaction of CD82-LEL with CADM1 Ig-like domain places the *trans*-interacting Ig1 domain of CADM1 close to the (*cis*) membrane instead of far (ca. 18 nm) from the membrane, preventing formation of a proper *trans* network [40, 39]. The two mechanisms are not mutually exclusive and can be achieved through ectodomain interactions, although involvement of membrane-helix interaction cannot be excluded based on our experiments.

Future research could reveal whether other members of the CADM family or the closely related CRTAM protein also interact with CD82 and whether other tetraspanins interact with CADM1, as tetraspanins can overlap in function [6, 54, 55, 56]. It is yet unclear whether CADM1 signaling via its intracellular PDZ domain or 4.1 binding motif is impacted by the interaction with CD82. Tetraspanins TSPAN7 and CD63 interact with their partner proteins via the PDZ domain and thereby block it for other proteins [57, 58]. A similar dual function on the extracellular and intracellular domain was shown for tetraspanin CD151 that regulates the conformation of the extracellular domain of integrins [59, 60] as well as integrin signaling [61]. Further research could also focus on the effect of post-translation modifications, especially N-glycosylation, on the complex. The discovery of CADM1 as an interaction partner of CD82, along with the inhibitory effect of CD82 on CADM1 oligomerization in *vitro* and in *cellulo*, opens new avenues for investigating the physiological functions of this interaction.

## Materials and Methods

### Constructs

Synthetic DNA encoding CD82 (UniProtKB: P27701.1, VAR_052326), CADM1 excluding the signal peptide (UniProtKB: Q9BY67.2), EGFR excluding the signal peptide (UniProtKB: P00533-1) and the linker used for CADM1-CD82 fusion protein (linker: AA-4xGGS-ENLYFQG-4xGGS-RS) was ordered from GeneArt. CD82 was ligated into expression vectors containing a C-terminal eGFP-StrepII_3_-tag or a C-terminal StrepII_3_-tag. CADM1 was cloned into expression vectors containing an N-terminal signal peptide and a C-terminal eGFP-StrepII_3_-tag, StrepII_3_-tag or His_6_-tag. EGFR was truncated after the transmembrane domain and amino acids 1-672 were ligated into an expression vector containing an N-terminal signal peptide and a C-terminal StrepII_3_-tag. Mutations and domain deletions of CADM1 were introduced using mutagenesis PCR. All expression vectors were provided by ImmunoPrecise Antibodies (Europe) BV.

### Small scale membrane protein purification

Expression vectors were transiently transfected into 4 mL of HEK293-E+ cells (ImmunoPrecise Antibodies (Europe) BV). Cells were harvested 4 days post-transfection by centrifugation (1,000 x g, 10 min, 4°C). Cell pellets were used fresh or flash frozen in liquid nitrogen and stored at -80°C. All purification steps were carried out at 4°C or on ice and with pre-cooled buffers. Cell membranes were solubilized with lysis buffer containing 150 mM NaCl, 25 mM HEPES pH 7.5, 1% w/v n-dodecyl-β-d-maltopyranoside (DDM) and EDTA-free Complete protease inhibitor tablets (Roche) with gentle mixing for 1 h. Cell lysates were centrifuged (100,000 x g, 45 min) and the supernatant was incubated with strep-tactin Sepharose® beads (GE Healthcare) for 2 h with gentle mixing. Beads were washed three times with ten column volumes of washing buffer containing 150 mM NaCl, 25 mM HEPES pH 7.5 and 0.025% w/v DDM. Protein was eluted using washing buffer supplemented with 3.5 mM d-desthiobiotin. Anti-His western blots were treated with a mixture of Monoclonal Anti-polyHistidine, Clone His-1 (Sigma) and Penta-His Antibody, Mouse anti-(H)5 (Qiagen) diluted 1:2,000 in blocking buffer (3% bovine serum albumin in PBS + 0.05% Tween-20) as primary antibodies. Anti-Flag western blots were treated with Anti-Flag M2 Monoclonal antibody (Sigma) as primary antibody. Anti-His and anti-flag western blots used 1:10,000 Goat anti-mouse IgG-HRP (BioRad) as secondary antibody.

### F-SEC and stability test

Samples for F-SEC were centrifuged at 20,000 x g, 4°C for 5 min. F-SEC was performed using a Superose 6 increase 10/300 GL column (GE Healthcare Life Sciences) or a Superdex 200 increase 10/300 GL column (GE Healthcare Life Sciences) connected to a Prominence UFLC system (Shimadzu) and running buffer containing 150 mM NaCl, 25 mM HEPES pH 7.5, 0.025% w/v DDM for membrane proteins and 200 mM NaCl, 25 mM HEPES pH 7.5 for CADM1(ECD)-His_6_ at a flowrate of 0.5 mL/min. The fluorescence of tryptophan (excitation: 275 nm, emission: 354 nm) or eGFP (excitation: 488 nm, emission: 509 nm) was detected by a RF-10AXL fluorescence detector (Shimadzu). Graphs were plotted using Origin 9.1 (OriginLab Corp.).

### Large scale membrane protein purification

For large scale expression in 200 mL, 1 L or 2 L HEK293-E+ cells, protein expression was carried out as described for small scale expressions. All steps were carried out at 4°C or on ice and with pre-cooled buffers. The cell pellet was washed once with PBS. Cell pellets were processed fresh or flash frozen in liquid nitrogen and stored at -80°C. The cells were lysed using buffer containing 150 mM NaCl, 25 mM HEPES pH 7.5, 10% v/v Glycerol, 10 µg/mL DNAse (Sigma Aldrich), 1% w/v DDM and EDTA-free Complete protease inhibitor tablets (Roche) with gentle mixing for 2 h. The lysate was centrifuged (100,000 x g, 45 min) and the supernatant was incubated with Strep-tactin Sepharose® beads (GE Healthcare) for 2 h with gentle mixing. Beads were washed with washing buffer, containing 150 mM NaCl, 25 mM HEPES pH 7.5 and 0.025% w/v DDM. Protein was eluted with 3.5 mM d-desthiobiotin supplemented washing buffer. For further purification, proteins were injected onto a Superdex 200 increase 10/300 GL or Superose 6 increase 10/300 GL column (GE Healthcare Life Sciences) equilibrated with washing buffer. Fractions with protein of interest were pooled and concentrated using a 100 kDa MWCO filter or 50 kDa MWCO (Amicon) for purifications of CD82-StrepII_3_ used for reconstitutions.

### Purification of CADM1(ECD) and CADM1(ECD, F42S)

Protein expression was carried out as described for membrane proteins. The DNA was transfected into 400 mL HEK293-E+ cells and cells were harvested six days post-transfection. The cells were pelleted twice by centrifugation (5,000 x g, 15 min, RT) and discarded. The supernatant was incubated with 3 mL Ni Sepharose excel beads (Cytiva), equilibrated with buffer containing 500 mM NaCl and 25 mM HEPES pH 7.5. Beads were collected into a gravity flow column and washed with 30 column volumes of buffer (500 mM NaCl, 25 mM HEPES pH 7.5, 25 mM imidazole). CADM1(ECD)-His_6_ or CADM1(ECD,F42S)-His_6_ were eluted in five fractions of 5 mL elution buffer (500 mM NaCl, 25 mM HEPES pH 7.5, 500 mM imidazole). Fractions containing protein were further purified by SEC using a Superose 6 increase column (Cytiva) equilibrated in running buffer (200 mM NaCl, 25 mM HEPES pH 7.5). Fractions containing pure protein were pooled and concentrated using 30 kDa MWCO filter (Amicon). Proteins were aliquoted, flash frozen in liquid nitrogen and stored at -80°C until further use.

### Mass spectrometry

Samples of CD82-eGFP-StrepII_3_ were expressed and purified as described in “Large scale purification of membrane proteins” and separated by 4-15% SDS PAGE. Gel bands were cut, fixed, destained, and digested with pig trypsin (Promega) following the procedure described elsewhere [62]. Resulting tryptic peptides were separated by liquid chromatography and analyzed by tandem mass spectrometry (LC-MS/MS) in a Q-Exactive Orbitrap Mass Spectrometer equipped with an Easy nLC1000 instrument (ThermoFisher) as also described in [62]. MS raw data files were analyzed using the MaxQuant software (v1.5.0.25) with the settings detailed elsewhere [63], except for the search against the Homo sapiens proteome retrieved from UniProt (01.01.2020) including known protein contaminants. Individual protein abundances were determined by label-free quantification (LFQ) and intensity-based absolute quantification (iBAQ) values.

### Size exclusion chromatography with multi-angle laser light scattering

SEC-MALLS was performed using samples diluted in running buffer and centrifuged at 20,000 x g for 5 min before injection onto a Superdex 200 increase 10/300 GL column (GE Healthcare Life Sciences) equilibrated in running buffer (200 mM NaCl, 25 mM HEPES 7.5) at a flow rate of 0.4 mL/min. Light scattering was measured using a miniDAWN TREOS multiangle light scattering detector (Wyatt), connected to a differential refractive index monitor (Shimadzu, RID-10A). Molecular weights were calculated using the ASTRA6 software (Wyatt) with an dn/dc value of 0.1745 mL/g considering six N-glycosylation sites in the extracellular domain of CADM1. The calibration was checked using 2 mg/mL Conalbumin (Sigma Aldrich) dissolved in running buffer at the start and end of the measurements. The molecular mass of Conalbumin was calculated using a dn/dc value of 0.185 mL/g. Graphs were plotted using Origin 9.1 (OriginLab Corp.).

### Liposome clustering assay

Dipalmitoylphosphatidylcholine (DOPC; Avanti) and 18:1 1,2-dioleoyl-sn-glycero-3-[(N-(5-amino-1-carboxypentyl)iminodiacetic acid)succinyl] (nickel salt) (DGS-NTA; Avanti) were mixed in chloroform at a molar ratio of 8:2. The chloroform was evaporated under a nitrogen flow while rotating to form a lipid film at the bottom of a rounded glass tube and further dried overnight under vacuum. The lipid films were rehydrated with 805 µL liposome buffer (150 mM KCl, 25 mM HEPES/KOH pH 7.5) resulting in calculated final concentrations of 3.17 mM DOPC and 0.79 mM of DGS-NTA for 1 h in a 37°C water bath and vortexed every 15 min. To completely dissolve the lipids, the mixture was freeze-thawed five times using liquid nitrogen for freezing and a 37°C water bath for thawing. The lipid mixtures were stored at -20°C until further use. Liposomes were thawed and extruded through an 800 nm filter. Liposomes used for reconstitutions were then additionally extruded through a 100 nm filter. To reconstitute freshly purified CD82-StrepII_3_ or EGFR(I-TM)-StrepII_3_, 90 µL of liposomes were incubated with DDM at a molar DDM:lipids ratio of 1:1 for 1 h while gently mixing at room temperature. CD82-StrepII_3_ was added to the liposomes at final concentrations of 7 µM and 3.5 µM and EGFR(I-TM) was added to a final concentration of 3.5 µM. Protein, detergent and liposomes mixtures were incubated at room temperature while gently mixing for 1 h. Detergent was removed stepwise by incubating three times with 20 mg of BioBeads (BioRad) equilibrated in liposome buffer for 30 min at RT and overnight with 100 mg at RT. Old BioBeads were removed in between the steps. Liposomes were mixed with CADM1(ECD)-His_6_ at different concentrations, 10 μL NanoTemper capillaries were dipped into solution until filled. Capillaries were measured ten times in technical duplicates for non-reconstituted liposomes and technical triplicates for reconstituted liposomes on a Prometheus Panta (NanoTemper).

### CRISPR/Cas9 mediated knock-out of CD82 in Jurkat

A guide RNA pair targeting the first coding exon of the human CD82 (GGATCACTGCGCCCAGGATC and GGGCTTCGGGGTGTGGATCC) was designed using the MIT CRISPR design tool [64] and cloned into the px335 Cas9 vector as described before [65]. Jurkat cells were transfected with 1.0 µg of each of the two gRNA plasmids of the pair and 0.5 µg pSGFP2-C1 using a Nucleofector 2b system (Amaxa, Lonza) according to the manufacturers guidelines using program T014 and Mirus nucleofection buffer (MirusBio). One day after nucleofection, GFP-positive cells were flow sorted on a FACSAria (BD Biosciences). After expanding the sorted culture, cells were stained for CD82 (A647-labeled, Clone NBP-2, Novus Biologicals) and CD82-negative cells were flow sorted on a FACSMelody (BD Biosciences).

### Airyscan microscopy

Jurkat T-cells were stained in suspension as previously described [66] for CD82 (B-L2, Novus Biologicals, 1/100) and CADM1 (3E1, MBL Life Science, 1/100). Secondary staining was performed with Alexa 647 goat anti-chicken (Thermo Scientific A-21449) (1/400) and Alexa 488 goat anti-mouse IgG1 (Thermo Scientific A21121) (1/400), respectively. Following staining, cells were adhered to poly-lysine-coated coverslips for 20 minutes at 4°C prior to fixing in 4% paraformaldehyde (PFA). Cells were then stained with DAPI for 10 minutes prior to mounting in Fluoromount G (SouthernBiotech 0100-01). Airyscan Microscopy was performed with a Zeiss LSM 900 microscope equipped with a 63 × EC Epiplan-NEOFLOUAR oil immersion objective. A Z-interval of 0.15 μm was used. Images were acquired and 3D rendering achieved using ZEN software. Images were subject to Airyscan processing following acquisition and were analyzed using ImageJ.

### Grid preparation and cryo-EM data collections

The complex of CADM1-His_6_ and CD82-StrepII_3_ was expressed in 2 x 1 L HEK293-E+ cells and purified as described. The 12 mL EV peak was plunge frozen at a concentration of 6.2 mg/mL and the 14 mL peak was plunge frozen at a concentration of 5.7 mg/mL on glow discharged R1.2/1.3 300 mesh Au holey carbon grids (Quantifoil). DOPC:DGS-NTA (8:2) liposomes were plunge frozen at lipid concentrations of 430 µM for liposomes coated with CADM1(ECD)-His_6_ on glow discharged R1.2/1.3 300 mesh Cu holey carbon grids (Quantifoil). All grids were plunge frozen in liquid ethane using a Vitrobot Mark IV system (Thermo fisher scientific) for 3.5 s at 8°C and 90% humidity. Grids were clipped and stored in liquid nitrogen until imaging. For the sample of CADM1-His_6_ and CD82-StrepII_3_ complex at 12 mL EV, 8,706 movies were collected on a 300 kV Krios1 microscope (Thermo fisher scientific) equipped with a K3 direct-detection camera (Gatan) using AFIS in EPU 2.20.0 at a magnification of 130,000 in super resolution mode resulting in a pixel size of 0.328 Å/px with a defocus range of -2.0 µm to -0.8 µm. Movies were collected with a total dose of 60 e-/Å^2^ distributed over 60 frames. For the CADM1-His_6_ and CD82-StrepII_3_ complex at 14 mL EV we collected a data set of 1,381 movies on a 200 kV Talos Arctica microscope (Thermo fisher scientific) equipped with a K2 direct-detection camera (Gatan) using AFIS in SerialEM at a magnification of 100,000 resulting in a pixel size of 1.36 Å/px with a defocus range of - 2.5 µm to -1 µm. Movies were collected with a total dose of 55 e-/Å^2^ distributed over 60 frames.

### Cryo-EM data processing

Data processing of CADM1-His_6_ and CD82-StrepII_3_ complex at 14 mL EV was carried out in CryoSPARC v3.3. For the data collected for the complex eluting at 12 mL EV most processing was carried out in CryoSPARC v3.3 and final refinement steps were done in CryoSPARC v4.5.3. Both data sets were corrected by Patch-based motion correction and CTF estimation and final exposures were manually curated. For the data set collected of the 14 mL complex peak, 1,221 micrographs were used for further processing. Initial particle picking was done with blob picker and a selection of 2D classes of particles showing one micelle and particles showing two micelles were used for template picking separately. Template picking was used for several rounds with increasing particle numbers until resulting 2D classes did not further improve in resolution and orientation of the particles. The final 2D classification of classes showing a single micelle included 235,125 particles and the final round for 2D classes showing two micelles included 152,173 particles. For the data set collected on the CADM1:CD82 complex eluting at 12 mL EV, 8,343 exposures were used for further processing. Initial particle picking was performed using blob picking followed by six rounds of particle cleaning in 2D. Particle cleaning in 3D was carried out by iterating ab initio modelling and heterogeneous refinements. 2D classes were created from the most promising 3D density model and used it for template picking, followed by three rounds of cleaning in 2D and iteration of ab initio modelling and heterogeneous refinements for cleaning in 3D. Particles selected based on 2D classification were used to train a TOPAZ model [67] for a final round of particle picking, resulting in a set of 1,349,274 particles. After one round of particle cleaning in 2D, we continued with particle cleaning in 3D as previously. An overview of the processing pipeline is illustrated in **Fig. S10**. To assess the density, an AlphaFold 3 prediction of CD82 and CADM1 was made [46] and CD82(LEL), CADM1(Ig1) and CD82 transmembrane domains and the SEL (CD82(TM, SEL)) were individually and manually fitted into the resulting densities using ChimeraX 1.8.

All graphs showing SEC, F-SEC profiles and bar graphs representing the calculated molecular weights from MALLS were created using Origin 9.1 (OriginLab Corp.). Images of Jurkat cells were processed using ImageJ. Figures displaying DLS data were created using PyCharm Community Edition 2024.2 (JetBrains). All figures containing protein densities were created using ChimeraX 1.8 (UCSF). Final figures were prepared using Illustrator (Adobe).

## Supporting information

EMDB54536 validation report

## Acknowledgements

We thank ImmunoPrecise Antibodies (Europe) BV for providing expression vectors and protein expression, the Protein Research Centre (Utrecht University) for providing the instrument for DLS measurements and J.W. Beugelink for support with SEC-MALLS. We thank the electron microscope centre (Utrecht University) and W. Noteborn from NeCEN (University Leiden) for data collections, M. Bergmeier and S.C. Howes for support during data collections at the electron microscope centre at Utrecht University and M. Vanevic for computational support during processing. A. Cabrera-Orefice and U. Brandt we want to thank for measuring mass spectrometry samples. For discussions and feedback on the manuscript we thank T.H.C. Brondijk.

## Data availability

Electron density maps of the CADM1:CD82 complex have been deposited in the EMDB with the ID EMD-54536.

## Supporting information

Supplemental figures S1-S12 [46, 47].

