## Supplemental Figures 1-12 for "Tetraspanin CD82 reduces the formation of CADM1 oligomers"

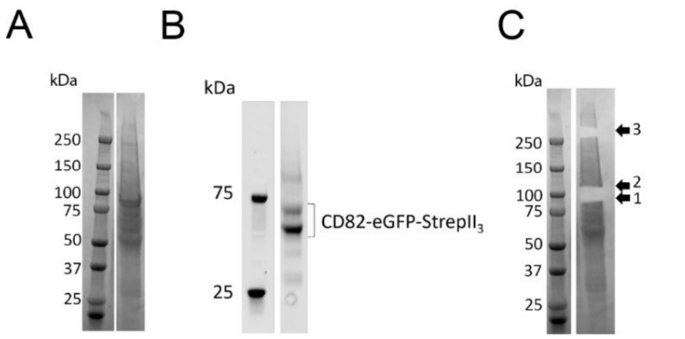


**Figure S1: Proteins co-purified with CD82-eGFP-StrepII$_3$** (A,B,C) 4-15% SDS-PAGE gel (A) stained with Coomassie, (B) eGFP signal and (C) after excising protein bands for mass-spectrometry.

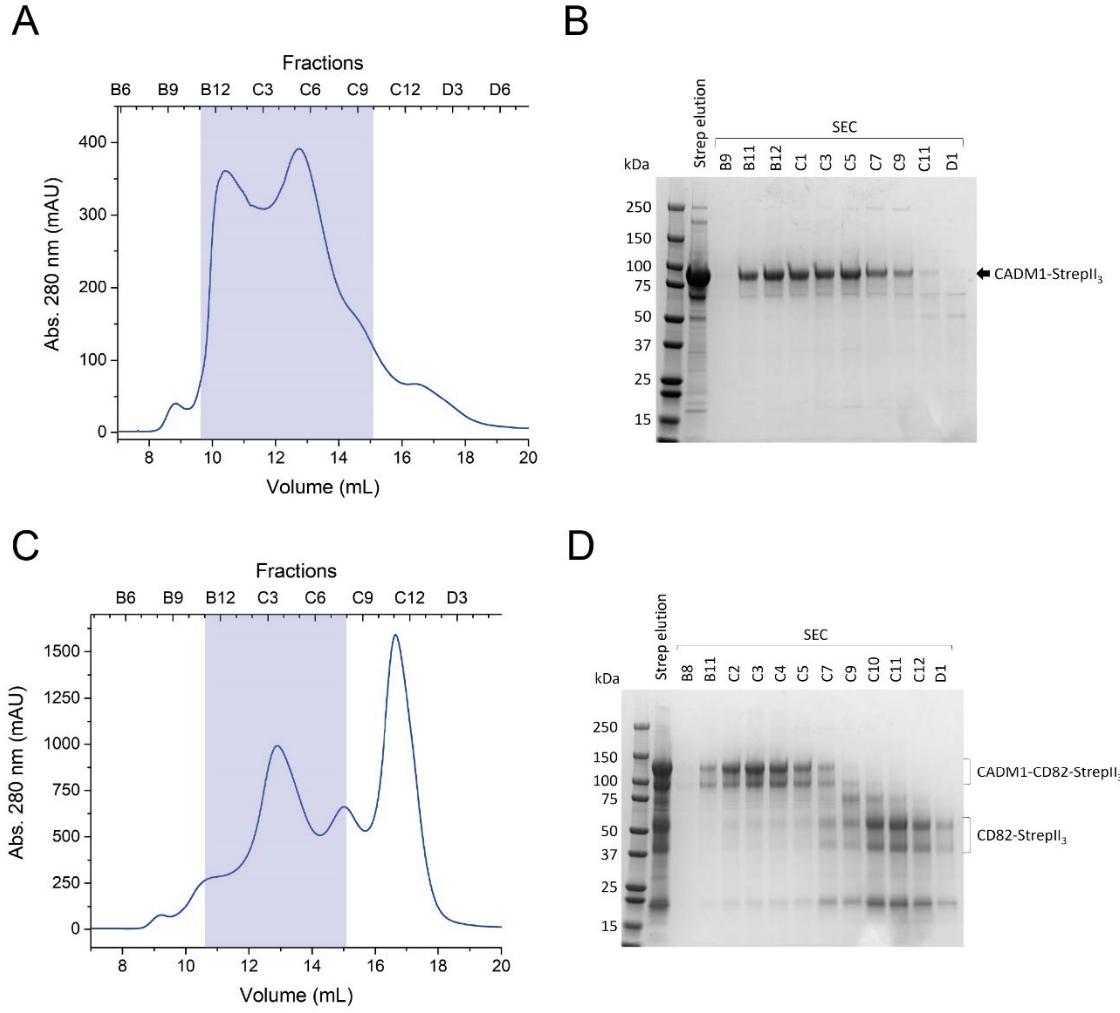


**Figure S2: Purification of CADM1-StrepII$_3$ and CADM1-CD82-StrepII$_3$.** (A) SEC of CADM1-StrepII$_3$ using a Superose 6 increase 10/300 GL column. Pooled fractions are marked with a translucent blue box. (B) 4-15% SDS-PAGE gel of single fractions of the SEC stained with Coomassie. (C) SEC of CADM1-CD82-StrepII$_3$ using a Superose 6 increase 10/300 GL column. Pooled fractions are marked with a translucent blue box. (D) 4-15% SDS-PAGE gel of single fractions of the SEC stained with Coomassie.

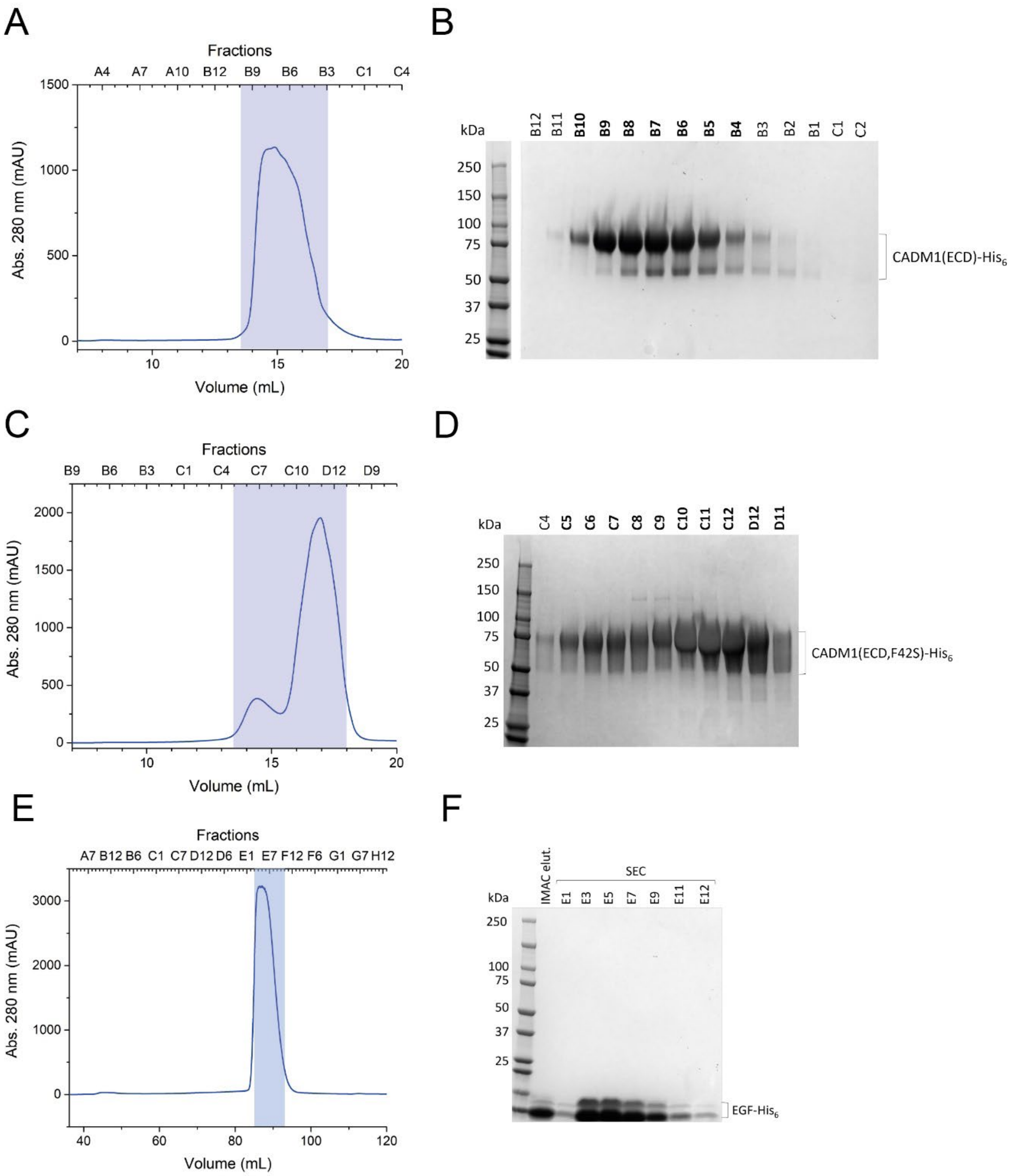


**Figure S3: Purification of CADM1(ECD)-His$_6$ and CADM1(ECD,F42S)-His$_6$.** (A) SEC of CADM1(ECD)-His$_6$ using a Superose 6 increase 10/300 GL column. Pooled fractions are marked with a translucent blue box. (B) Individual fractions were analysed by 4-15% SDS-PAGE stained with Coomassie. Pooled fractions are labelled in bold. (C) SEC of CADM1(ECD,F42S)-His$_6$ as in panel A. (D) Individual fractions were analysed by 4-15% SDS-PAGE stained with Coomassie.

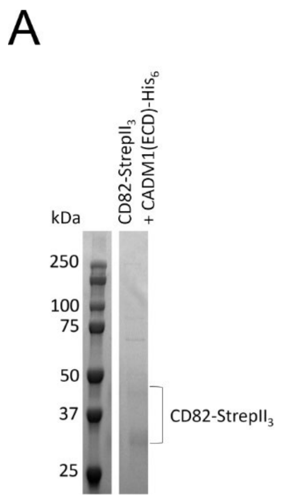


**Figure S4: Interaction between CADM1(ECD)-His$_6$ and CD82-StrepII$_3$ interaction site.** (A) 10% SDS-PAGE gel stained with Coomassie and, (B), anti-His western blot of streptavidin purified CD82-StrepII$_3$ with 50 µg (0.1 mg/mL final concentration) CADM1(ECD)-His$_6$ added to the detergent solubilized cell lysate after ultracentrifugation.

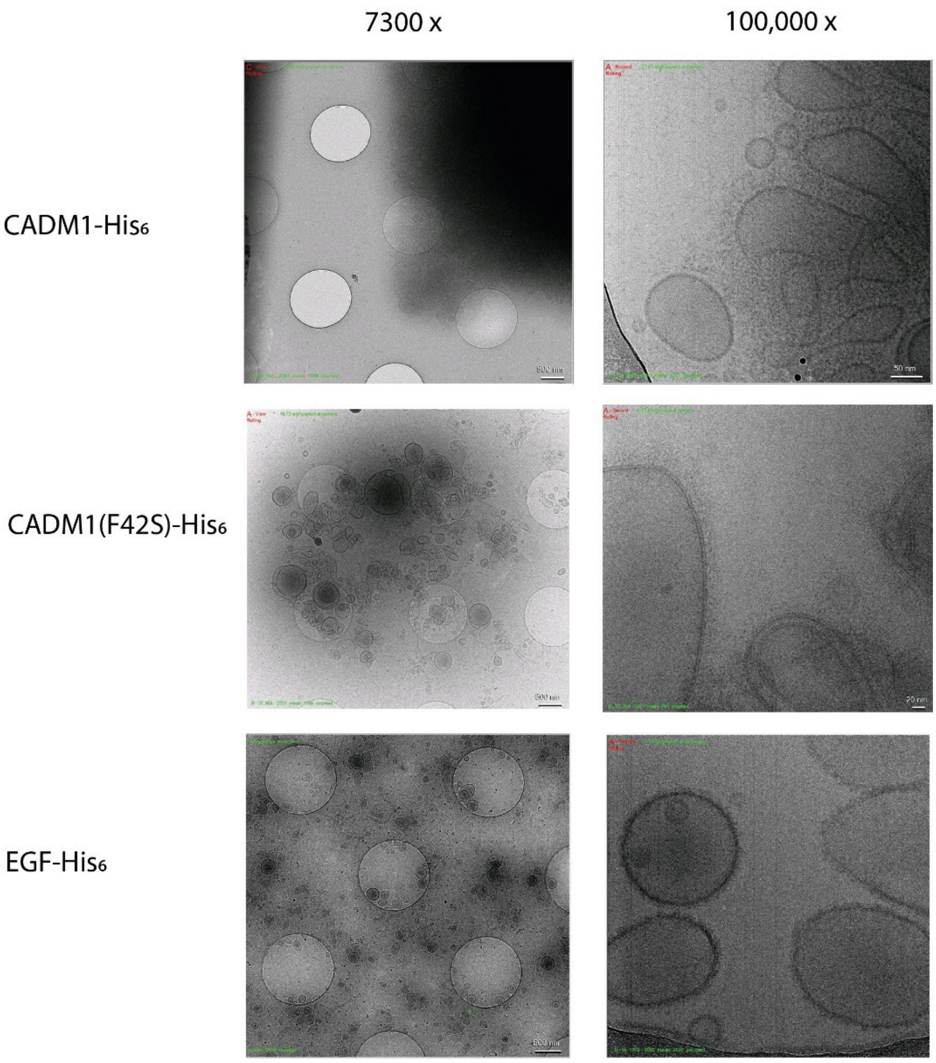


**Figure S5: Liposomes coated with different proteins.** Liposomes were coated as indicated in the figure and pictures were taken at magnifications of 7300x and 100,000x using a Talos Arctica electron microscope (200 kV).

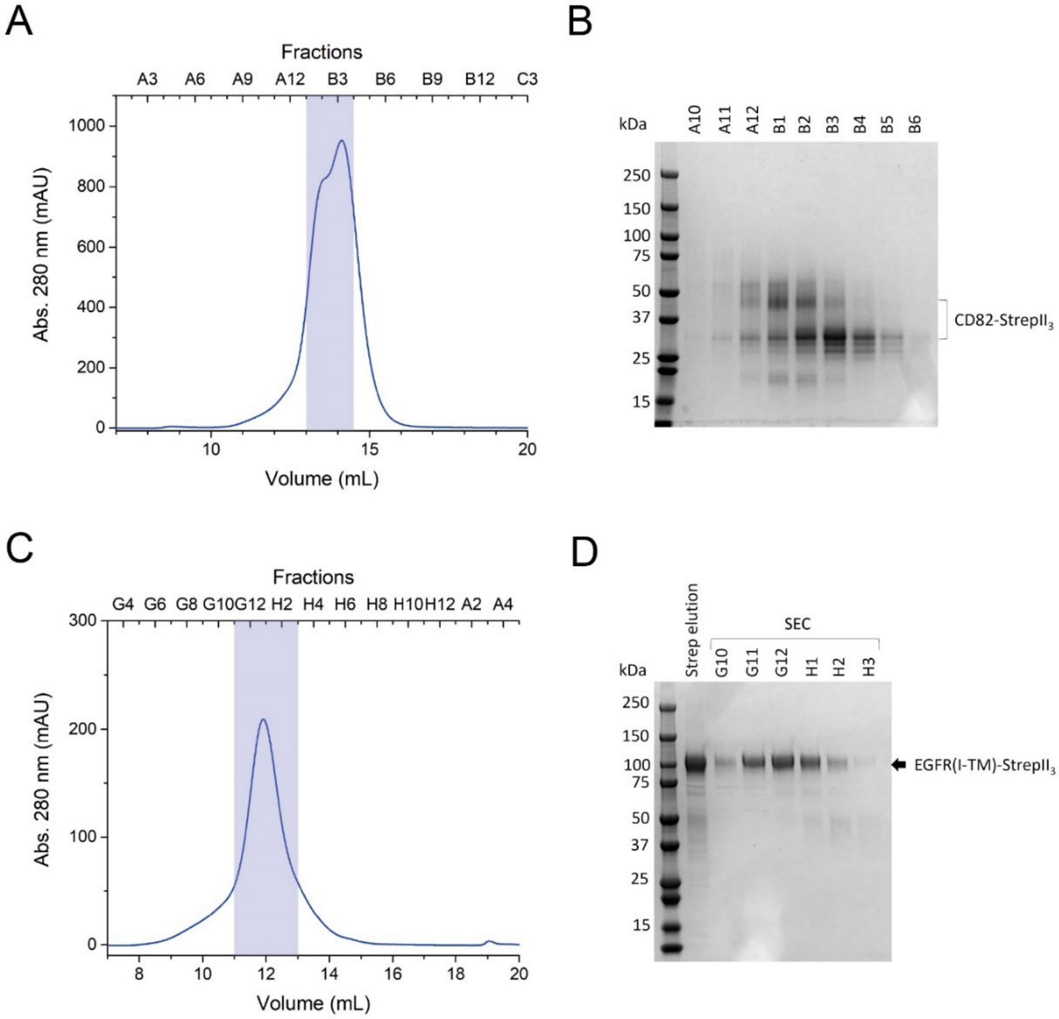


**Figure S6: Purification of CD82-StrepII$_3$ and EGFR(I-TM)-StrepII$_3$ for reconstitution in liposomes.** (A) SEC of CD82-StrepII$_3$ using a Superdex 200 increase 10/300 GL column. Pooled fractions are marked with a translucent blue box. (B) 4-15% SDS-PAGE gel of single fractions of the SEC. (C) SEC of EGFR(I-TM)-StrepII$_3$ using a Superdex 200 increase 10/300 GL column. Pooled fractions are marked with a translucent blue box. (D) 4-15% SDS-PAGE gel of single fractions stained with Coomassie.

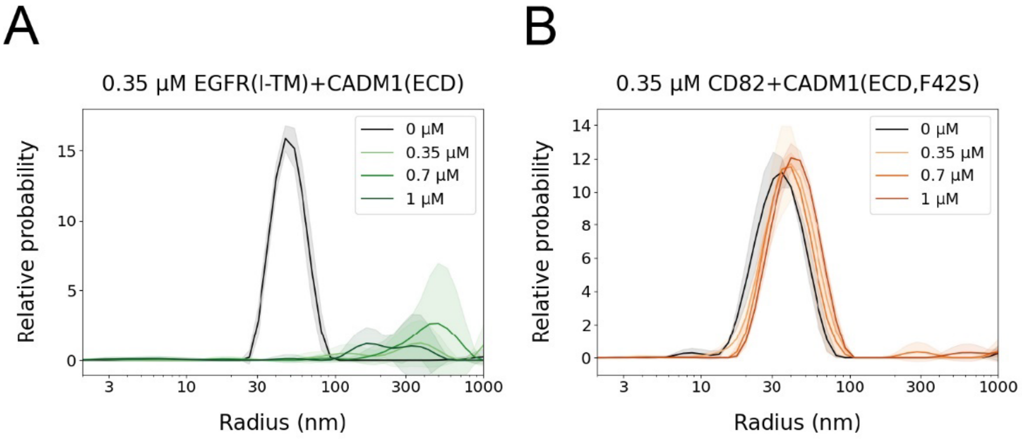


**Figure S7: Controls of liposome clustering assay.** (A,B) DLS by DOPC:DGS-NTA (8:2) liposomes (A) reconstituted with (A) EGFR(I-TM)-StrepII$_3$ and coated with CADM1(ECD)-His$_6$ and (B) reconstituted with CD82-StrepII$_3$ and coated with CADM1(ECD,F42S)-His$_6$. The same liposome concentrations were used as in Fig. 2C-E.

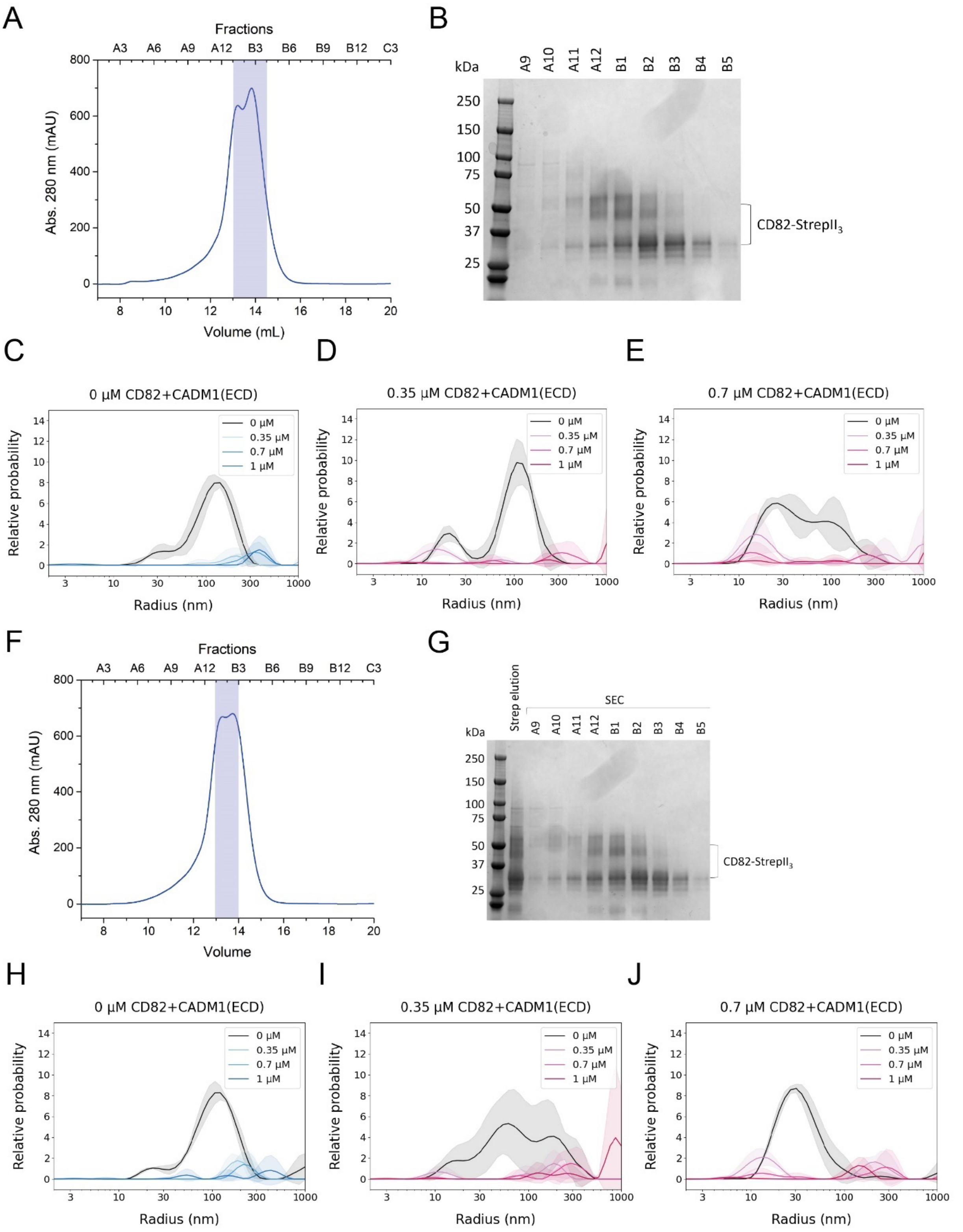


**Figure S8: Repetitions of liposome clustering on assay.** (A) SEC of CD82-StrepII$_3$ using a Superdex 200 increase 10/300 GL column. Used fractions are marked with a translucent blue box. (B) 4-15% SDS gel of individual SEC fractions stained with Coomassie. (C-E) DLS by DOPC:DGS-NTA (8:2) liposomes reconstituted with different concentrations of CD82-StrepII$_3$ and each coated with different concentrations of CADM1(ECD)-His$_6$. (F)-(J) display a repetition of the same experiments as in panel A-E.

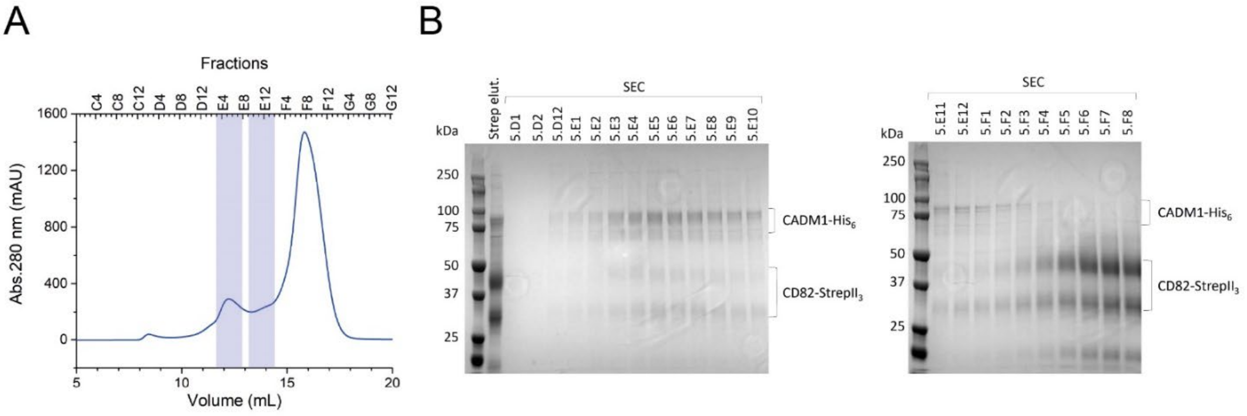


**Figure S9: Purification of CADM1-CD82 complex for cryo EM.** (A) SEC profile of streptavidin-purified CADM1-$His_6$ and CD82-$StrepII_3$ on a Superose 6 10/300 GL column. Analysed fractions are marked with a blue transparent box. Fractions of each box were combined into two separate pools. (B) Coomassie-stained 10% SDS-PAGE gels of sample injected onto the SEC column (Strep elut.) and individual SEC fractions.

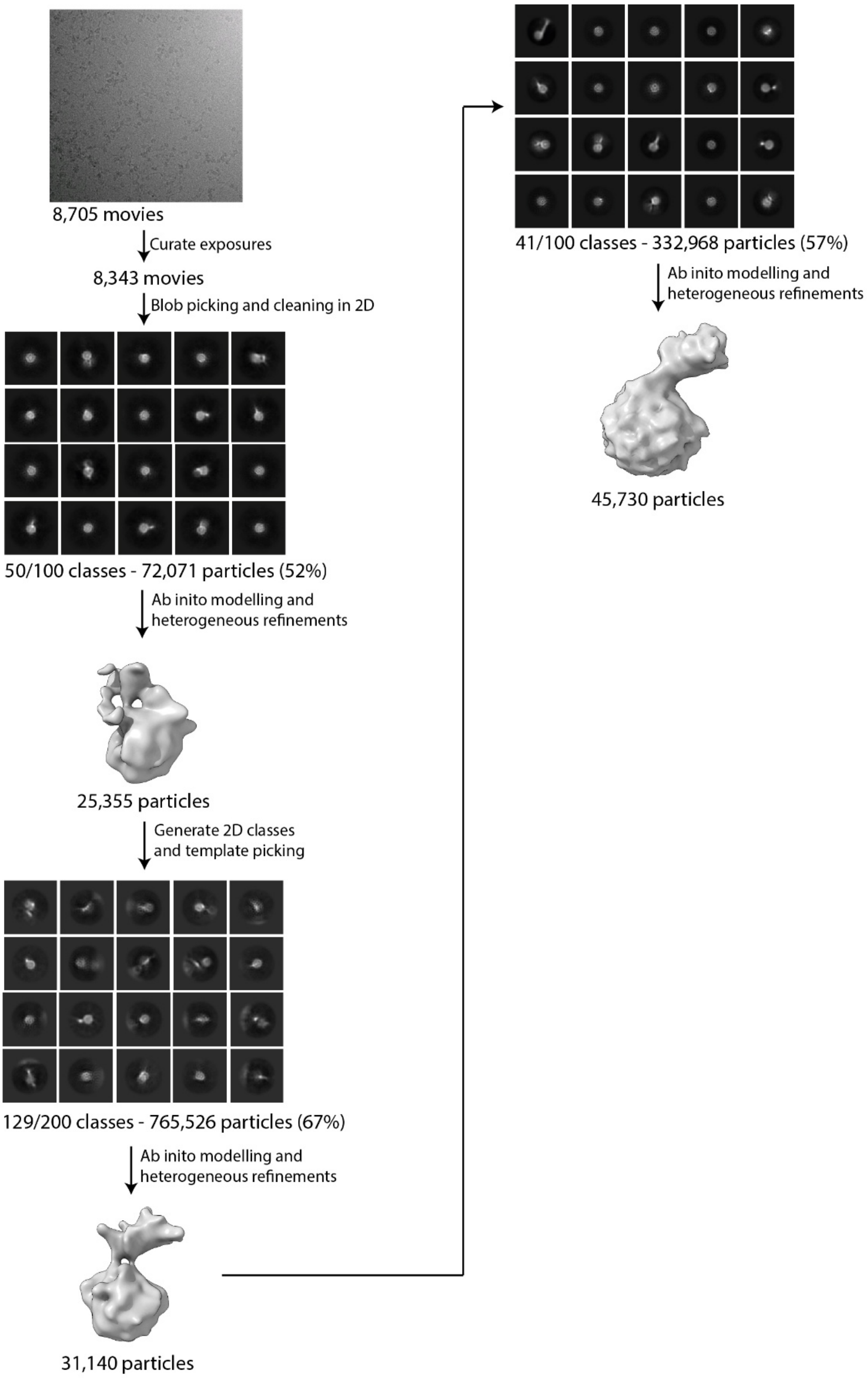


**Figure S10: Processing pipeline of CADM1-CD82 complex.** Processing steps leading to the density shown in figure 6. Only a selection of representative 20 2D classes are shown. Ellipses indicate the target of local refinements.

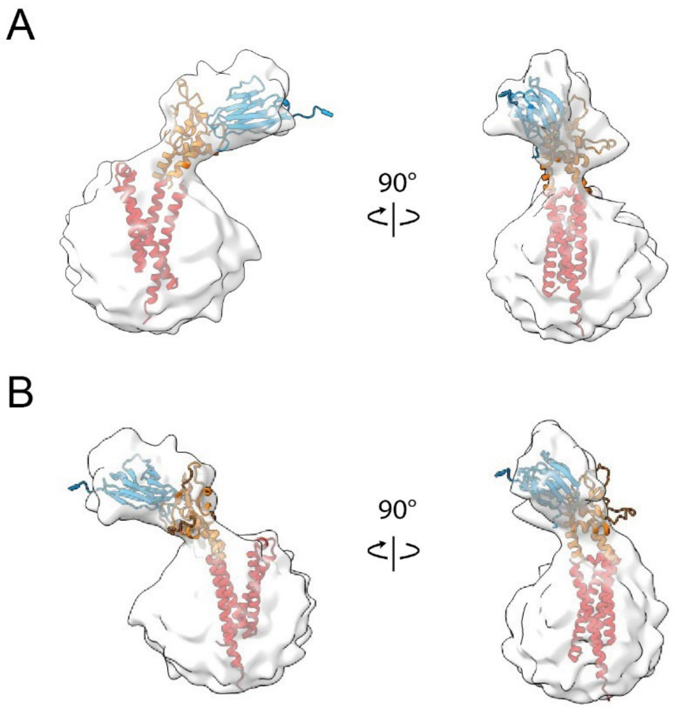


**Fig S11: Determination of handedness.** (A) Volume shown in Fig. 5C as front view and side view. (B) Density of Fig. 5C flipped over y-axis. Rigid bodies from an AlphaFold3 prediction were fitted in the volume as shown in Fig. 5C.

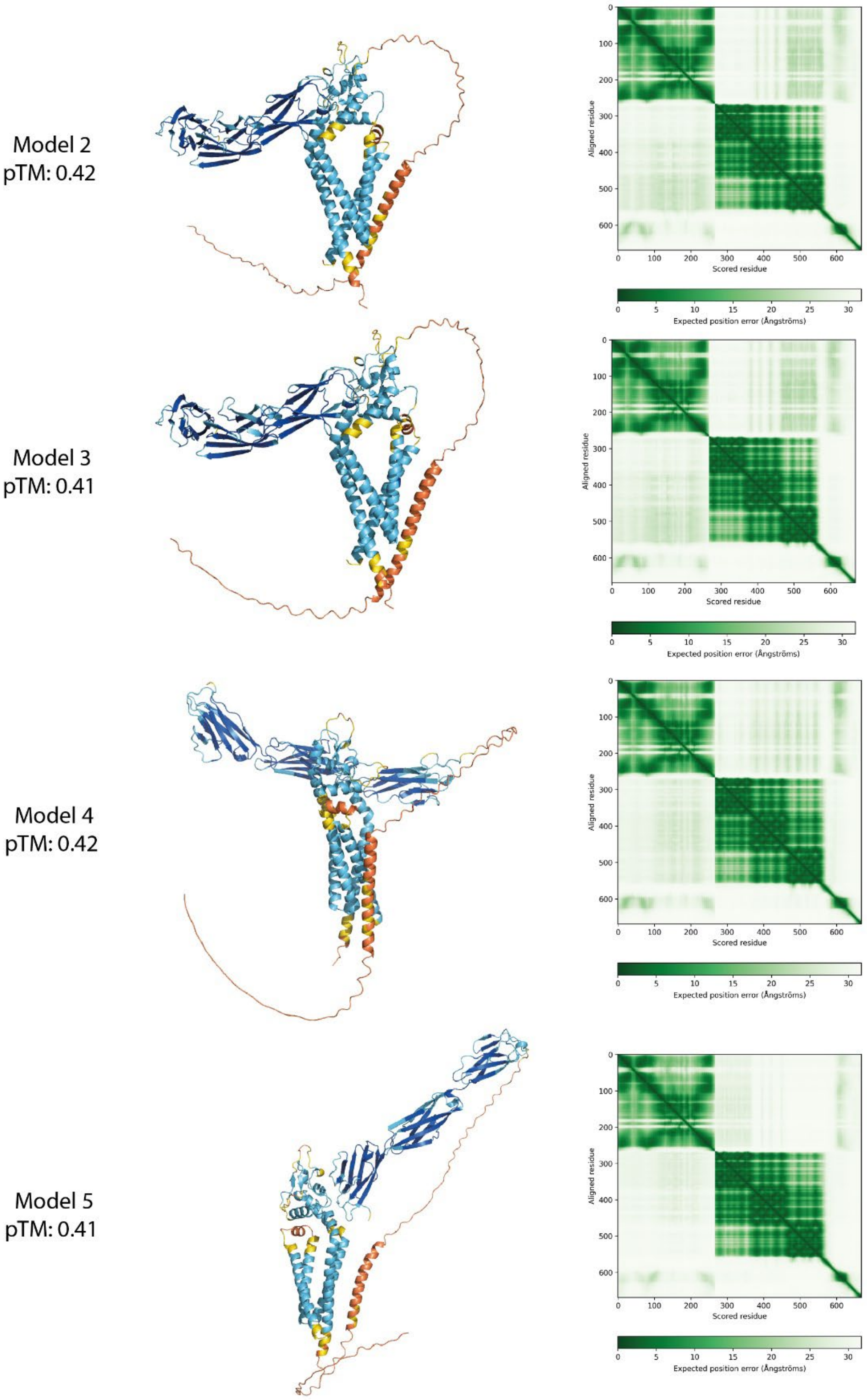


**Figure S12: AlphaFold3 prediction of CADM1-CD82 complex.** AlphaFold3 prediction of additional four models with pTM scores as indicated in the figure colored with AlphaFold pLDDT scores and predicted aligned error matrix [46, 47].
